# Light and electron microscopy continuum-resolution imaging of 3D cell cultures

**DOI:** 10.1101/2021.07.02.450855

**Authors:** Edoardo D’Imprima, Marta Garcia Montero, Sylwia Gawrzak, Paolo Ronchi, Ievgeniia Zagoriy, Yannick Schwab, Martin Jechlinger, Julia Mahamid

## Abstract

3D cell cultures, in particular organoids, are emerging models to investigate healthy or diseased tissues. Understanding the complex cellular sociology in organoids requires integration of imaging modalities across spatial and temporal scales. We present a multi-scale imaging approach that traverses millimeter-scale live-cell light microscopy to nano-scale volume electron microscopy by performing 3D cell cultures in a single carrier amenable to all imaging steps. This allows to follow organoids growth, probe their morphology with fluorescent markers, identify areas of interest and analyze their 3D ultrastructure. We demonstrate this workflow on mouse and human 3D cultures, and use automated image segmentation to annotate and quantitatively analyze subcellular structures in patient-derived colorectal cancer organoids. Our analyses reveal local organization of diffraction-limited cell junctions in compact and polarized epithelia. The continuum resolution imaging pipeline is thus suited to foster basic and translational organoid research by simultaneously exploiting the advantages of light and electron microscopy.

**Highlights:**

- Establishment of 3D cell cultures in sample carriers directly amenable to high-pressure freezing (HPF)
- 3D cell cultures in HPF carriers allow drug treatment and live-cell imaging
- Multi-scale imaging of 3D cultures from live-cell light microscopy to volume electron microscopy
- Establishments of HPF conditions for mouse and patient-derived organoids
- Deep-learning automatic segmentation of ultrastructural detail and quantitative data-mining reveal different subcellular organization associated with epithelium polarity

## Introduction

Cell culture systems are indispensable for a wide range of basic and pre-clinical studies. When modelling tissue-specific processes or multifaceted diseases like cancer, conventional two-dimensional (2D) cell cultures largely fail to recapitulate the complexity and cellular heterogeneity of the normal and cancerous tissue (Baker and Chen, 2012). Conversely, three-dimensional (3D) cell cultures can deliver more accurate representations of cell-cell and cell-extracellular matrix (ECM) interactions, cell communication (Kretzschmar and Clevers, 2016) and cell division (Knouse et al., 2018). 3D cancer culture models allow studying tumor microenvironment, cell heterogeneity, and cell invasion (Kapałczyńska et al., 2016; Zanoni et al., 2020), as well as the effect of anti-cancer drugs (Lancaster and Huch, 2019). Thus, patient or cell line-derived 3D culture systems have the potential to bridge the gap between simplified 2D models and organismal models that are more expensive and inaccessible to many imaging or high-throughput methods. 3D cultures can be established by anchorage-independent cell suspensions, scaffold-based approaches (including embedding in natural laminin rich hydrogels (termed Matrigel, (Hughes et al., 2010)) or synthetic ECM), tissue slices, or air-liquid interfaces (Koledova, 2017; Shamir and Ewald, 2014). Commonly, 3D cultures are derived from immortalized cell lines, induced pluripotent stem cells (iPSC), embryonic stem cells (ESCs), or primary cells dissociated from animal or human tissues. When grown in 3D, immortalized cell lines can form hollow spheres or solid spheroids depending on their respective epithelial polarization phenotypes. Organoids are derived from stem cells or primary cells from healthy or diseased tissues and reflect the respective original tissue organization in 3D culture (Fatehullah et al., 2016; Simian and Bissell, 2017).

Imaging is a powerful tool to study the complex 3D cellular sociology in organoids. Bright-field microscopy enables long-term imaging of organoid development at a micrometer scale with minimal phototoxicity (Hof et al., 2021). Confocal and light-sheet microscopy permit the study of specific cellular processes at higher resolution either using fluorescent reporters for cell types, organelles, or proteins (Day and Davidson, 2009), or cell-permeable dyes (Ettinger and Wittmann, 2014). However, the inherent opaque nature of large organoid cultures (> 200 µm) requires optical clearing (Tomer et al., 2014). Thus, to ensure effective fluorescent labelling and imaging, organoids are commonly extracted from the Matrigel (Dekkers et al., 2019), preventing post clearing image correlation. Further, immune-based fluorescence necessitates sample permeabilization for precise tagging. Both approaches require chemical fixation fixation that precludes live-cell imaging (Ertürk et al., 2012). To visualize ultrastructural detail at the nanometer scale, researchers generally rely on 2D imaging of ultrathin sections by transmission electron microscopy (TEM) (Knott and Genoud, 2013). Such data are often associated with low throughput, sectioning artifacts and limited field of view on the specimen. The biggest limitation of this technique, when imaging 3D cell cultures, is the lack of volumetric information which is essential to thoroughly appreciate cellular organisation in space. Conversely, volume electron microscopy (Peddie et al., 2022) and in particular focused ion beam-scanning electron microscopy (FIB-SEM) is an ideal solution to this problem. This method involves progressive specimen slicing by FIB surface ablation, followed by SEM imaging (Giannuzzi and Stevie, 2005; Heymann et al., 2006). FIB-SEM thus generates a series of images that can cover a relatively wide field of view at nanometer-scale resolution and extensive z-slicing yields relatively large volumes of 3D ultrastructural information (Heymann et al., 2006; Narayan and Subramaniam, 2015).

However, none of these imaging methods alone can provide a complete structural and functional understanding across spatial scales of cells in the context of the complex multicellular environment of a 3D organoid. To alleviate such shortcomings, correlative approaches that combine the power of light and electron-based imaging modalities have been developed (Ganeva and Kukulski, 2020; Kukulski et al., 2011; Peddie et al., 2022). Yet, their implementation in organoid research has remained limited due to the diverse requirements these imaging modes have, which are rarely met by one single specimen preparation; established electron microscopy (EM) specimen preparation requires sample crosslinking by chemical fixation at room temperature, leading to disruption of the cells’ fine ultrastructure (Mollenhauer, 1991). Cryogenic fixation instead uses low temperatures to stabilize the sample under physiological conditions, i.e. fully hydrated. Freezing must retain the water molecules in an amorphous solid state in a process called vitrification (Dubochet and McDowall, 1981) and for larger objects, including 3D cell cultures, vitrification is achievable by high-pressure freezing (HPF) (Moor, 1987). Here, samples that are up to 200 microns thick are sandwiched between two metal disks, known as planchets or HPF carriers. Then, the specimen protected between the metal carriers is pressurized to 2000 bar, immediately followed by cooling with liquid nitrogen within less than 200 milliseconds. For room temperature EM, the cryo-fixed samples are then subjected to freeze-substitution and stained with heavy metals to provide contrast in EM imaging (Kellenberger, 1987). A variation of this procedure reduces the amount of metals in the freeze-substitution cocktail to only 0.1% uranyl acetate, allowing preservation of endogenous fluorescent signal for targeting of specific regions of interest (ROI) in large sample volumes by fluorescence imaging after embedding (Kukulski et al., 2011; Nixon et al., 2009), while still providing enough contrast for FIB-SEM imaging (Ronchi et al., 2021). Such preparation methods are easily applicable for specimens that are amenable to manual handling for cryo-fixation, including small model organisms or dissected tissues. However, the fragility of 3D organoids grown in soft matrices precludes such manipulation.

It is therefore beneficial to perform the 3D cell cultures and live imaging in sample carriers that are directly amenable to cryo-fixation. Here, we show that it is possible to perform 3D cell culture in common HPF carriers coated with a biocompatible metal, and develop a seamless workflow to image the cultures from their initial development to high-resolution FIB-SEM. This establishes a multiscale imaging pipeline in which the same organoid is tracked across the different modalities and scales. We apply the pipeline to mouse mammary gland organoids, human breast cancer spheroids, and patient-derived colorectal cancer organoids. For the latter, we capitalized on deep learning algorithms to automatically segment and mine over 10^5^ cubic microns of FIB-SEM data to gain ultrastructural insight into the heterogenous distribution of subcellular structures that cannot be resolved by diffraction-limited light microscopy. Based on the quantitative analysis of the large-scale segmentations, we show that microvilli and cell junctions concentrate locally to seal off lumina in organoid regions with denser cell-packing. We thus demonstrate advantages of continuum resolution imaging on fragile 3D organoids with complex and heterogenous morphologies, when cultured in a single specimen carrier.

## Results

### Establishing a multiscale imaging pipeline

In multimodal imaging pipelines, EM has the most stringent requirements for specimen preparation due to its high resolving power. HPF is the only approach to preserve the native ultrastructure of multicellular samples. We tested whether common gold-coated copper HPF carriers as readily available supports are compatible with long-term 3D cell culture and the subsequent imaging modalities (Fig 1A). We used 200 μm deep HPF carriers featuring a nominal 0.62 mm^3^ sample volume and 3.14 mm^2^ surface area that is exposed to the culture medium. This results in a surface-to-volume ratio that is 2.5 times larger than our standard conditions, namely 3D cell culture gels (90 µl Matrigel drop) in tissue culture (TC) dishes. We used the latter according to published protocols (Jechlinger, 2015) and for culturing patient-derived organoids. We performed the 3D cell culture directly in HPF carriers by pipetting in the recess ∼1 μl of cell suspension mixed with Matrigel (Materials and Methods). We then placed the carriers in multi-well TC-dishes supplied with the culture medium. To aid handling and to prevent floating of the small carriers, we fixed the carriers with a droplet of Matrigel to 18 mm glass coverslips. With this setup, we could directly monitor the 3D cell culture growth by stereo-microscopy up to 24 days and correlate it with subsequent live-cell confocal imaging (Fig. 1A, B). Because our aim was to probe organoid morphology for EM, we only collected stereo microscopy images at single time points to monitor the organoid growth. Nevertheless, acquisition of time-series of confocal fluorescence microscopy (4D imaging) is technically possible using this setup. Next, we performed high-pressure freezing at chosen time points of the culture. Freeze-substitution, heavy metal staining that retains fluorescence signal in the specimen and Lowicryl HM20 resin embedding were carried out in preparation for room temperature FIB-SEM (Ronchi et al., 2021). The resin block was then imaged by confocal fluorescence microscopy to relocate features previously identified in the specimen by live-cell imaging and define regions of interest (ROI). We generated landmarks on the resin block surface by two-photon laser ablation to facilitate identification of the ROIs and guide their high-resolution imaging in subsequent FIB-SEM (Ronchi et al., 2021). We then acquired FIB-SEM data of the targeted areas, resulting in an imaging pipeline covering all spatial scales from 10^−3^ to 10^−9^ m on the same organoids. The pipeline may require up to 10 days, with the freeze-substitution being the longest step (Fig 1B). In summary, transferring fragile organoids into HPF carriers is a cumbersome procedure which requires organoids removal from the ECM (Triffo et al., 2008); this may alter their morphology and prevents image registration pre and post cryo-fixation. Thus, by establishing 3D cultures directly in HPF carriers, sample handling and all the imaging steps require minimal manual intervention.

**Figure 1.**
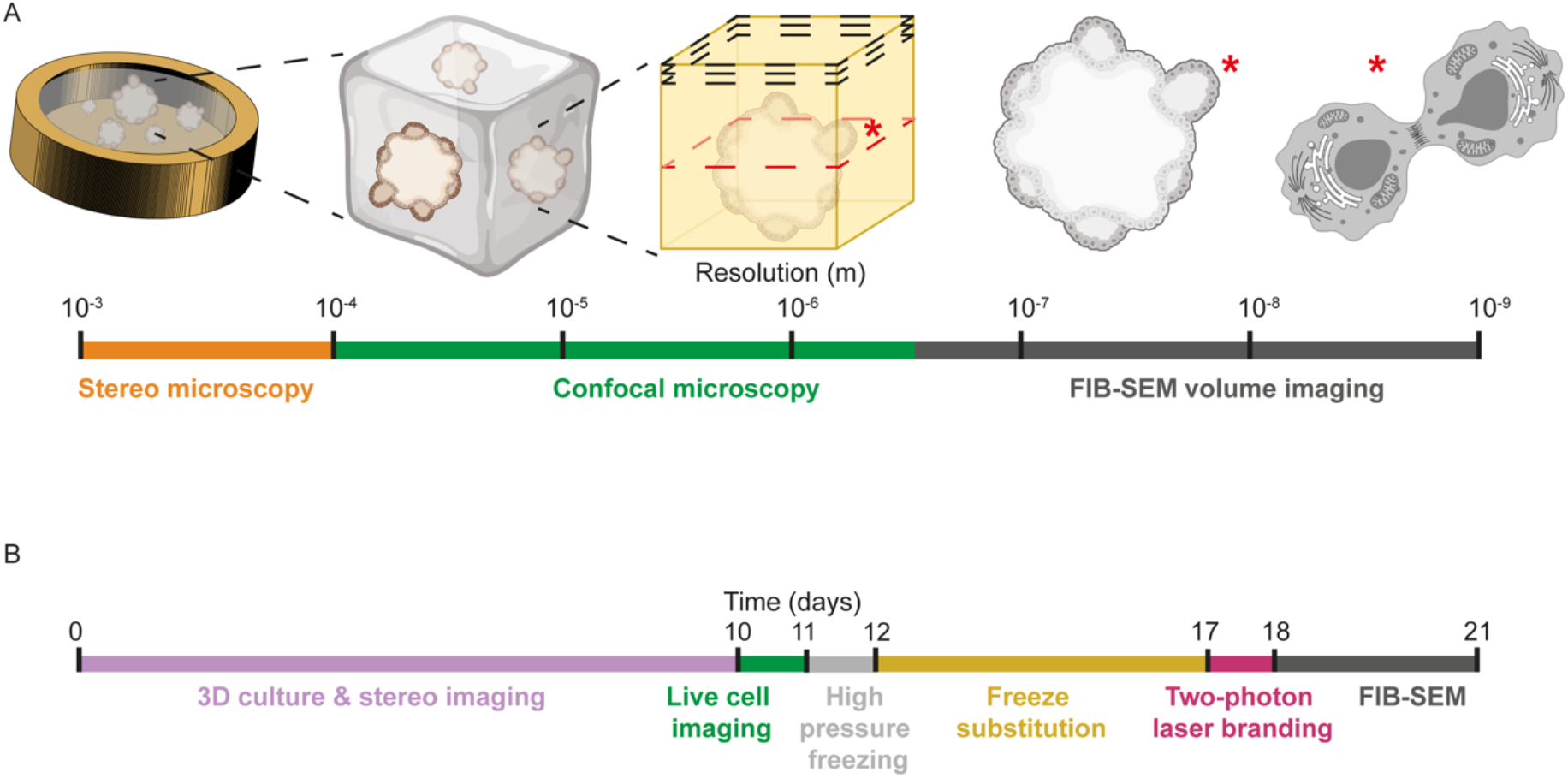
3D cell cultures in HPF carriers enable correlative imaging from the millimeter to the nanometer scale. **A**, a multiscale imaging pipeline of 3D cell cultures in HPF carriers encompasses millimeter-scale stereo light microscopy, 3D confocal fluorescence microscopy prior to and following cryo-fixation, and nanometer-scale FIB-SEM volume imaging. Asterisks indicate single cells that can be targeted and followed throughout the imaging pipeline. **B**, timeline of 3D cell culture for room temperature sample preparation and imaging.

### HPF carriers are compatible with diverse 3D cell culture models

To validate the compatibility of the developed pipeline for organoid research, we examined three different types of 3D cell cultures: models of primary healthy and tumorigenic (under doxycycline oncogene induction) epithelia derived from mouse mammary glands, patient-derived human colorectal cancer, and a human breast cancer cell line (Fig 2 and S1). We grew our model of healthy mammary gland organoids from single-cell suspensions derived from transgenic mice expressing H2B-mCherry as a nuclear fluorescent marker. Within 14 days, mammary gland organoids grown in the HPF carriers displayed the expected single layer of epithelial cells arranged around a lumen, similar to standard culture conditions (Fig 2A, B and S1A,B) (Alladin et al., 2020). The fluorescent signal of the transgenic H2B-mCherry reporter allowed us to inspect organoids distribution and morphology over several days of cell culture (Fig S1E). We performed immunofluorescent staining for the apical-basal cell polarity marker epithelial Cadherin (E-CAD) and the tight-junction protein Zonula occludens-1 (ZO-1), which localized in the apical cell membrane, on organoids cultured in HPF carriers and TC dishes. These showed that the lumen is lined by the apical membrane of the polarized epithelial cells, while the basal side faces the ECM in both HPF carriers (Fig 2B) and TC (Fig. S1B), confirming that characteristics of polarized epithelia are preserved. In addition, immunostaining of tissue-specific differentiation markers on fixed samples directly in the HPF carriers showed that the organoids are predominantly composed of luminal mammary epithelial cells expressing Keratin-8 (K8) and a few basal/myoepithelial mammary cells expressing Keratin-14 (K14) (Fig 2C). Moreover, we verified that activation of c-Myc oncogene expression by doxycycline addition in the HPF carriers lead to the development of more solid tumorigenic organoids, reminiscent of the tumor-induced organoids grown on the dishes (Alladin et al., 2020) (Fig S1H-K).

**Figure 2.**
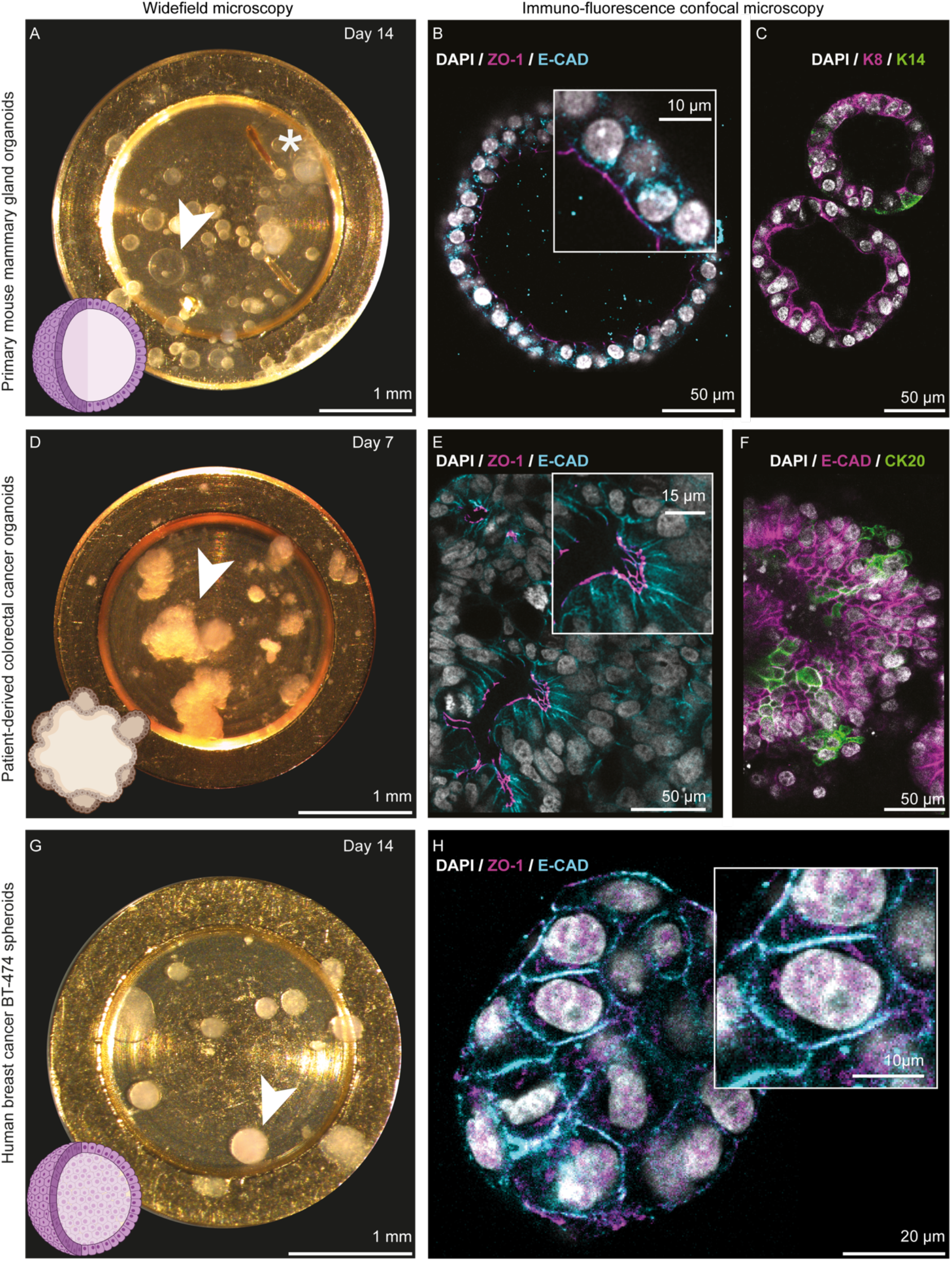
Preservation of organoid-specific 3D multicellular morphologies in HPF carriers. Mouse mammary gland organoids growth in HPF carriers (**A**) at the indicated time point of the culture (also shown in Fig S1E). Arrowhead indicates a single organoid. Asterisk indicates marks that can be applied to the bottom of the HPF carriers with blunt-tip tweezers prior to Matrigel deposition to assist imaging by different light microscopes. Bottom left: schematic of luminal organoid morphology. **B**, immunostaining of polarity markers: the cell-adhesion protein epithelial cadherin (E-CAD, cyan), tight junctions protein Zonula occludens-1 (ZO-1, magenta). Cell nuclei stained with DAPI (white). Inset: details of cell-cell and cell-lumen interfaces. **C**, immunostaining with tissue-specific markers: luminal Keratin-8 (K8, magenta) and basal/myoepithelial Keratin-14 (K14, green). Cell nuclei stained with DAPI (white). **D**, patient-derived colorectal cancer organoids growth at the indicated time point of the culture (also shown in Fig S1F). Arrowhead indicates a single organoid. Bottom left: schematic of organoid morphology. **E**, immunostaining of polarity markers E-cadherin (E-CAD, cyan), tight junctions (ZO-1, magenta). Cell nuclei stained with DAPI (white). Inset: details of cell-cell and cell-lumen interfaces. **F**, immunostaining with tissue-specific markers: E-cadherin (E-CAD) and Cytokeratin-20 (CK20, green), a colon cell marker; cell nuclei stained with DAPI (white). **G**, human breast cancer spheroids growth at the indicated time point (also shown in Fig S1G). Arrowhead indicates a single organoid. Bottom left: schematic of solid spheroid morphology. **H**, immunostaining of tissue-specific markers: E-cadherin (E-CAD, cyan), tight junctions (ZO-1, magenta). Cell nuclei stained with DAPI (white). Insets: details of cell-cell interfaces. See also Fig S1.

Patient-derived colorectal cancer organoids grow within a week as irregularly-shaped clusters (Fig 2D). These organoids display similar morphology when grown in HPF carriers and in standard 3D gels on TC dishes (Fig 2D, Fig S1C). Fluorescence live-cell confocal imaging of the patient-derived organoids in HPF carriers stained with nuclear dye Hoechst-33342 showed multiple cell layers with small lumina (Fig S1F). Immunofluorescent staining in both HPF carriers (Fig 2E) and TC dishes (Fig S1D) for E-CAD and ZO-1 showed that the expected apical-basal cell polarity is recapitulated in both culture conditions. Concurrently, the organoids expressed E-CAD and the Cytokeratin-20 (CK20) marker specific for colon tissue (Kummar et al., 2002) (Fig 2F).

Human breast cancer cells BT-474 are a cell line which develops as regularly-shaped spheroids when embedded in Matrigel and cultured in HPF carriers for up to 14 days (Fig 2G). Confocal live-cell imaging confirmed that these spheroids form as a solid sphere without lumina in HPF carriers (Fig S1G) and do not display apical-basal cell polarity when probed by immunofluorescent staining of E-CAD and ZO-1 (Fig 2H) (Florian et al., 2019).

These results demonstrate that luminal mouse mammary gland organoids, patient-derived colorectal cancer organoids and human breast cancer spheroids grown on HPF carriers display morphologies, cellular composition, and cell polarity reminiscent of organoids grown on TC dishes. Therefore, HPF carriers are applicable for a wide range of 3D cell cultures, and are compatible with confocal microscopy performed on live and fixed organoid samples as the first step in a multiscale imaging pipeline.

### Small molecule live dyes for correlative imaging of 3D cell cultures

Genetically engineered cell lines or animal models are frequently used to prepare 3D cell cultures, allowing the introduction of fusion proteins as fluorescent reporters. The production of transgenic reporters is time consuming, and is impractical for speedy translational molecular medicine in the case of patient-derived organoids, although recent publications show promising results in this direction (Shimokawa et al., 2017). Alternatively, live dyes can come as a time saving solution. In figure 2, we demonstrated the broad applicability of organoid cell culture in HPF carriers on organoids derived from transgenic mice with or without fluorescent reporters, and with two widely studied human-derived systems that lack genetically encoded fluorescence. For the latter, we investigated the possibility to used cell-permeable fluorescent live dyes to visualize cellular structures by confocal microscopy. However, small-molecule live dyes are often incompatible with many fixation procedures. Specifically, they are washed off by organic solvents during chemical fixation and dehydration required for EM preparations, unless they are supplied during the embedding step (Biel et al., 2003) which, in turn, prevents correlation with pre-embedding live-cell imaging. Nevertheless, mass spectrometry studies showed that live dyes may be resilient to the freeze substitution (Pfeiffer et al., 2000). Thus, we tested four common live dyes with different chemical and physical properties: Hoechsts-33342, SiR-actin, FM4-64 and BODIPY 493/503. In all 3D cell culture systems, we used a combination of Hoechst-33342 to mark cell nuclei and one of the 3 different dyes. All live dyes infiltrated organoids grown in HPF carriers, allowing live imaging of cell nuclei, of SiR-actin stained cortex in human BT474 spheroids (Fig 3, Fig S2 A-C) and in mammary gland organoids (Fig S2 D-F), of BODIPY 493/503 stained lipid droplets in mammary gland tumor organoids (Fig S2 G-I), and of cell membranes stained with FM4-64 in patient-derived colorectal cancer organoids (Fig S2 J-L). Furthermore, the live dyes were largely preserved in the gentle freeze-substitution procedure used here. In agreement with their chemical properties, hydrophilic dyes were not affected by freeze substitution (Hoechsts-33342, SiR-actin, Fig 3B, C) allowing detection of fluorescence before and after freeze substitution. The fluorescence signal of the membrane marker FM4-64 appeared diffuse after freeze-substitution, but permitted discerning cell boundaries to some extent (Fig 3D, E). The hydrophobic lipid droplet marker BODIPY 493/503 was completely removed during the washing step with organic solvents in freeze substitution (Fig 3F). Conversely, HPF alone did not affect the fluorescence signal of lipophilic live dyes as determined by direct imaging of different HPF samples with a cryo-confocal microscope (Fig 3G-I). For cryo-confocal imaging, we found the Leica HC PL FLUOTAR 50x 0.80 Numerical Aperture (NA) objective to provide the best compromise for high NA and long working distance (1.02 mm) to account for HPF carriers’ depth (> 100 µm) and large size of the organoids (> 100 µm). Thus, live dyes present an alternative for genetic tagging in fluorescence-based correlative imaging in 3D cultures.

**Figure 3.**
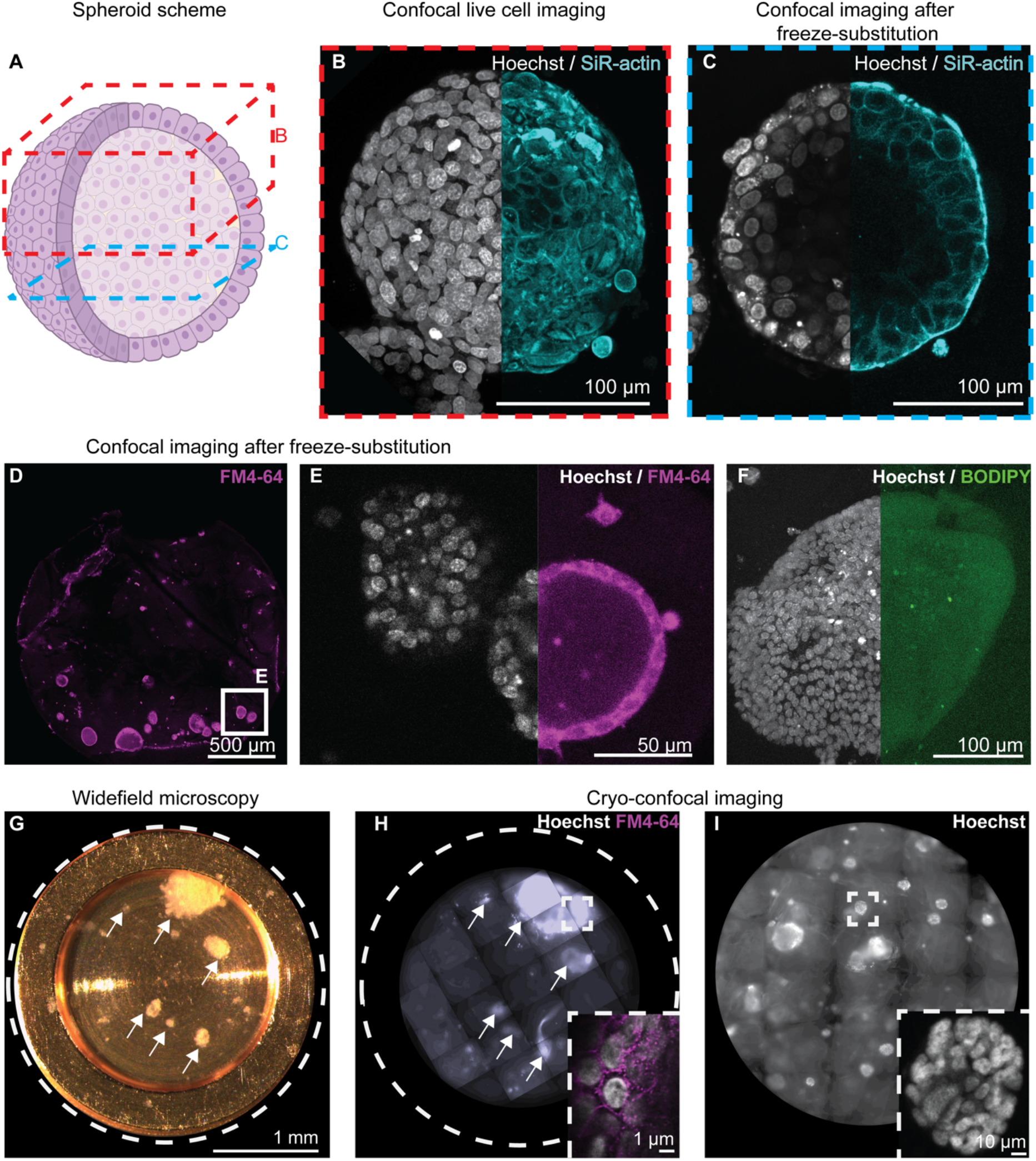
Preservation of fluorescent live dyes in cryo-fixation and freeze-substitution. **A**, schematic representation of a BT474 human breast cancer spheroid. Locations of confocal imaging before (red, B) and after (blue, C) freezing and freeze-substitution are indicated. **B**, maximum intensity projection of the live spheroid confocal volume stained with live dyes for detection of cell nuclei (Hoechst-33342, grayscale) and F-actin (SiR-actin, cyan). **C**, a confocal plane of the exposed resin block surface. **D-F**, Confocal imaging of primary mouse mammary gland organoids stained with membrane and nuclear live dyes (FM4-64 in magenta, Hoechsts-33342 in grayscale, respectively) after freeze-substitution. **F**, Tumor-induced mouse organoid stained with nuclear (Hoechst-33342, greyscale) and lipid-droplet (BODIPY 497/503, green) live dyes. **G**, light microscopy of patient-derived colorectal cancer organoids (white arrows) cultured in HPF carriers before high-pressure freezing. **H**, Maximum intensity projection tiled scan light microscopy at cryogenic temperature of the carrier in (**G**) after high-pressure freezing. Inset of an organoid in (**H**) with nuclear (Hoechst-33342, grayscale) and membrane (FM4-64, magenta) live dyes. **I**, light microscopy at cryogenic temperature of BT474 human breast cancer spheroids cultured in HPF carriers. Inset, an organoid in (**I**) with nuclear (Hoechst-33342, grayscale) live dye.

### FIB-SEM volume imaging of 3D organoids

Following localization of ROIs in the resin block by confocal imaging and generation of surface landmarks by two-photon laser branding in the same microscope, we performed FIB-SEM volume imaging on selected organoids. It is well known that effective vitrification by HPF varies from sample to sample. The embedding ECM or medium, its water content, sample thickness, and even the metabolic state of different cells within the same specimen affect the quality of vitrification (Sitte et al., 1987). Ice crystals growth can induce organelles collapse, segregation of compartments and aggregation of macromolecules. The EM data showed that organoids display a different freezing behaviour from their embedding medium, the Matrigel mix, which reveals fiber-like features (Fig 4A, B). Fully grown mammary gland organoids developed in 6-8 days in culture, during which cells polarized and self-organize in monolayered acini. Before reaching this stage, cells are intertwined in several layers within crowded acini (lumen up to 15% of the whole organoid volume) that show high cell packing density (Fig 4A). FIB-SEM data of organoid at this stage of growth generally showed good ultrastructural preservation with fine details of centrioles, mitochondria and condensed chromatin (Fig 4C). Conversely, in mature monolayer organoids, the lumen comprises the majority (up to 60%) of the organoid volume (Fig 2B, 4D). In this scenario, the polarized cells displayed typical freeze substitution artifacts including cracks and membrane detachment at cell-cell and cell-ECM interfaces (Fig 4D, E) due to resin polymerization-induced shrinkage. Also, freezing damage is evident in nuclei and in the cytosol (Fig 4E). Here, dehydrated biological material segregates to form high contrast, branched structures which grow from water crystal nucleation points and spread through the surrounding sample.

**Figure 4.**
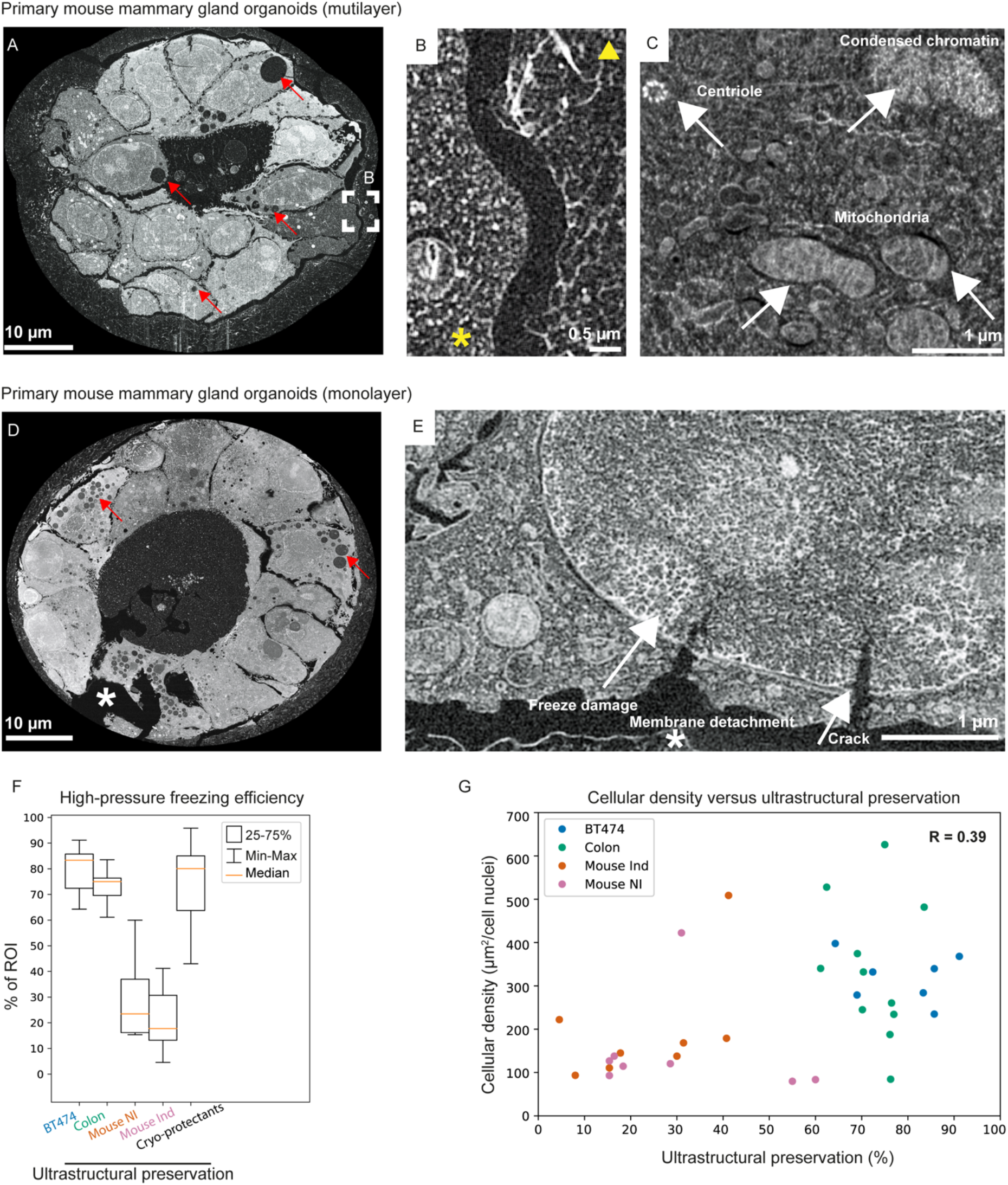
Assessment of structural preservation of 3D cell cultures. **A-C**, example of FIB-SEM slice of multilayer mouse organoids after seven days of culture. **B**, details of the cytoplasm characteristic freezing (left, asterisk) versus Matrigel (right, arrowhead). **C**, a well-preserved cell with condensed chromatin, a centriole, and mitochondria. **D-E**, monolayer organoids show freezing damage and extensive cracks, with membrane detachment. Red arrows in **A** and **D** highlight cell vacuoles characteristic of mouse mammary gland organoids. **F-G**, quantification of HPF efficiency by thin sections TEM and its correlation with cell density in organoid (G) for the four 3D culture types in the absence of cryo-protectants: human breast spheroids (BT474), human colorectal cancer organoids (Colon), doxycycline-induced tumorigenic (Mouse Ind), healthy (Mouse NI) mouse mammary gland organoids that were additionally supplemented with cryo-protectants (Cryo-protectants).

Despite the inherent heterogeneity of freezing quality expected in HPF, our preliminary observation of early multi-layered versus late monolayered mammary gland organoids suggested that the preservation of the cellular ultrastructure improved with increased local density of cells within the organoids and reduced size of the lumen. To provide a semi-quantitative description of the freezing quality, we prepared thin sections from the HPF 3D cultures for TEM imaging, allowing us to image larger areas of the different organoids and to probe diverse and distant regions across the HPF carriers. We prepared thin sections from doxycycline-induced tumorigenic (Mouse Ind), and healthy (Mouse NI) mouse mammary gland organoids, human spheroids (BT474) and patient-derived colorectal cancer organoids. The sections were cut orthogonal to the HPF carrier surface aiming for organoids located in the bulk of the Matrigel (Fig S3A). We performed TEM imaging from six to ten organoids per sample, spaced between 90 to 400 µm in the direction of cutting progression from the edge of the HPF carrier towards its center (Fig S3A). Although limited to 2D sampling, TEM thin sections have the advantage of higher resolution and the possibility to survey larger areas, compared to FIB-SEM, by tiling multiple imaged spots (Fig S3B). We visually inspected each tile for the presence of freeze damage artifacts and scored the quality of structural preservation. Overall, our survey shows that compact organoids like human colorectal cancer organoids achieve high ultrastructural preservation, with a minimum of 60% of structurally intact regions and median of 83%, or human breast cancer spheroids with a minimum 60% and median 75% (Fig 4F). Conversely, we obtained less consistent success for mouse organoids with large lumina varying between 23% and 17% of median ultrastructural preservation (Fig 4F). More specifically, human colorectal cancer organoids grow as large and compact structures that protrude into the Matrigel with multiple branches featuring one lumen (Fig S3C). Unlike the mouse organoids, human colorectal organoids show better preserved cell ultrastructure and a larger number of cells with little or no freezing damage (Fig S3B, Fig 4F). The human breast cancer spheroid represented the densest cell system that is devoid of a lumen and exhibited minimal freezing damage. Freezing damage was restricted to the center of the sample, as expected from theoretical considerations of heat transfer during HPF (Fig S4, Fig 4F). Structural damage was further confined to cell nuclei, and the occasional cytosolic segregation in the TEM section was clearly restricted to one or two cell layers at the spheroid center (Fig S4). Finally, we estimated the cellular packing density for all three systems as the ratio between the organoids total area divided by the number of cells, approximated by counting the nuclei in the TEM sections. Our data suggest that more compact organoids suffer less freeze damage despite the use of an identical embedding matrix and carrier dimensions (Fig 4G).

Optimization of HPF conditions is generally required for each specimen type (Möbius et al., 2010). This typically consists of embedding or infiltrating the sample with release agents like 1-hexadecene or lecithin, and cryo-protectants, such as, glycerol, dimethyl sulfoxide, bovine serum albumin, yeast paste, fish gelatin, polyvinylpyrrolidone, Dextran, sucrose, and Ficoll. These compounds work by directly inhibiting the nucleation of ice crystals or increasing the sample cooling rates by suppressing the heat released in the crystallization process (McDonald et al., 2007). We therefore sought to improve the vitrification of the least preserved 3D culture sample encountered, namely the mammary gland mouse organoids, by introducing cryo-protectants prior to cryo-fixation. We found that dipping the HPF carriers for 1 minute before HPF in Cellbanker 2, a common medium for preserving cells in liquid nitrogen (see material and methods section for details), or in 20% Ficoll (70.000 MW) dissolved in cell culture medium, improved the sample quality after HPF (Fig S5). Quantification of the ultrastructural preservation as described above showed improvement from ∼30% to ∼77 % (median values, Fig 4F).

In summary, Matrigel-embedded organoids are challenging samples for successful HPF because of the high-water content within the embedding medium, that inevitably causes freeze damage artifacts. On the other hand, Matrigel provides the necessary physiological environment for 3D cell cultures. We found that organoids with local high density cell packing are more resilient against freeze damage and provide adequate ultrastructural preservation. Moreover, we show that different organoid systems require optimisation in preparation for HPF, which can be achieved by careful selection of additives.

### Multimodal imaging of 3D organoids with continuum spatial resolution

Using the HPF carriers as a container for 3D cell culture allows minimal manual intervention during their development and facilitates correlation of diverse modalities to achieve continuum resolution imaging from millimeter to nanometer scale. We demonstrate the full imaging pipeline by performing 3D cell culture of human breast cancer spheroids directly in the HPF carriers and monitoring their growth by stereo microscopy (Fig 5A). At a chosen development stage of the culture, each carrier was stained with live dyes and transferred into 35 mm glass-bottom dishes for the acquisition of live-cell confocal volumes (Fig 5B). This provided information on the overall architecture of the 3D cell culture and allowed for selection of cells within specific organoids for higher resolution imaging. Subsequently, the samples were high-pressure frozen and processed with a gentle freeze-substitution protocol to preserve the fluorescence signal (Fig 5C). Two-photon laser surface branding of the resin-embedded samples at selected positions identified by fluorescence aided the targeting of FIB-SEM data with micrometer precision (Fig 5D). We identified the surface branded targets in the SEM, performed metal deposition and trench preparation following established protocols (Fig 5 E-H) and acquired FIB-SEM data with isotropic sampling of 10 nm^3^/voxel (Movie 1). Typically, precise correlation of light and electron microscopy (CLEM) data in 3D still requires tedious manual positioning and registration. Alternatively, dedicated hardware that integrates a widefield light microscope into a dual beam FIB-SEM provide means for direct registration of the data (Smeets et al., 2021). However, because we performed all imaging steps on the same specimen without manual intervention, we could precisely overlay the FIB-SEM volume data of just a few cells within the ∼ 7*10^6^ µm^3^ volume of the fluorescence data (Fig 5 I-K). The durability of the FIB ion source and the required lengthy milling time limit FIB-SEM imaging to 50-60 µm depth from the specimen surface. However, one could easily access any organoid within the HPF carrier by removing the excess of resin before the FIB-SEM acquisition by ultramicrotomy guided by the preserved fluorescence after freeze-substitution. Finally, we show that the FIB-SEM data acquired from six cells within the BT474 organoids with our protocols were of sufficient contrast and quality by segmenting the nuclei and mitochondria in the cells targeted for acquisition (Fig 5 L-N).

**Figure 5.**
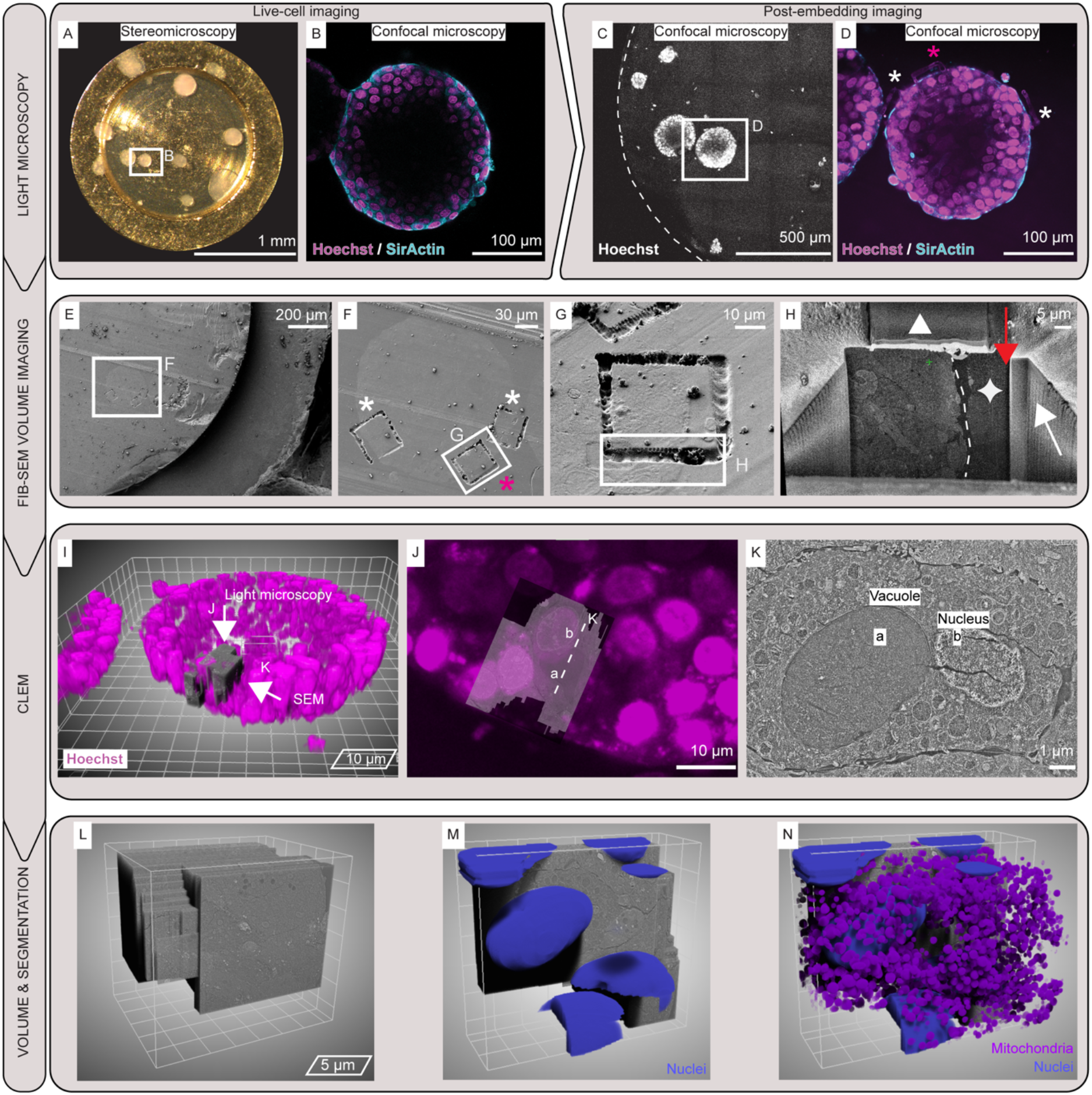
Correlative imaging of 3D cell cultures from the millimeter to the nanometer scale. **A**, stereomicroscopy of BT474 human breast cancer spheroid cultured in HPF carrier. **B**, live cell confocal slice of the indicated spheroid in **A** showing cell nuclei (Hoechst-33342, magenta) and F-actin (SiR-actin, cyan). **C**, maximum intensity projection tiled scan of the specimen from **A** after HPF and freeze-substitution. Frame indicates the spheroid imaged in **B**, and enlarged in **D. D**, confocal plane after freeze substitution. Surface laser brandings were introduced at positions of interest (white asterisks) to guide FIB-SEM volume imaging (magenta asterisk). Cell nuclei (Hoechst-33342, magenta) and F-actin (SiR-actin, cyan). **E**, SEM view of the sample in **D. F**, brandings (white asterisks) and FIB-SEM target (magenta asterisk) before (**G**) and after trench milling (**H**) showing the organoid outer edge (white dash) next to the embedding Matrigel (white star). Details of the acquired volume are shown in Movie 1. The white triangle indicates the protective platinum layer above the milling volume. Red and white arrows indicate the milling direction and the progression, respectively. **I**, correlated light and electron volume imaging of targeted cells from the organoid in **H** (magenta asterisk in **D** and **F**). CLEM overlay of the nuclear fluorescence (Hoechst, magenta) of the organoid in **B**-**D** (mesh size 10 µm) and the targeted FIB-SEM acquisition in **H** (grey). Top arrow indicates the fluorescence imaging direction shown in J. Side arrow indicates the imaging direction in the FIB-SEM acquisition shown in K. Post embedding fluorescence and SEM imaging planes are displayed in **J** and **K**, respectively, where the correlated location of a vacuole (a) and nucleus (a) are highlighted. **L**, 3D representation (mesh size 5 µm) of the FIB-SEM acquisition, and automatic segmentation of cell nuclei (**M**), combined with mitochondria (**N**).

### Characterizing ultrastructural heterogeneity of tumorigenic organoids

Intratumor heterogeneity is a prominent feature of cancer and a major impediment to personalized medicine (Almendro et al., 2013). Thus, we focused the imaging and analysis on our model of patient-derived colorectal cancer organoids. This organoid system exhibited high level of structural preservation by high-pressure freezing (Fig S3). Fluorescence microscopy showed that fully grown organoids exhibited regions with different cell packing arrangements depending on the presence of branched lumina (Fig 2D-E, Fig S2L). We took advantage of this heterogenous morphology to investigate whether and how it translates to structures at the subcellular level. We used nuclear staining (Hoechst-33342) to define three positions that differed in their tissue architectures for FIB-SEM acquisitions (Fig S6A, B). From those, we collected datasets of 121, 113 and 204 µm^3^ volumes at a resolution of 15×15×20 (x,y,z) nm^3^/voxel over 72 hours per dataset (Fig S6A-D). This represents the most extensive 3D ultrastructural study of organoids to date. In this imaging modality, colorectal cancer organoids display complex networks of cells, which could not be captured in its entirety by either 2D TEM or light microscopy alone: heterogenous cell packing spaced by multiple lumina (Movie 2) alternated with polarized parts of the organoids (Movie 3 and Fig S6E) or with highly packed epithelium morphology (Movie 4). This 3D volume imaging provided the opportunity to capture cancer-related events, such as entotic cells, that are rarely observed either due to insufficient resolution or absence of specific fluorescent markers (Fig S6F) (Fais and Overholtzer, 2018). In this non-apoptotic process, some cells internalize adjacent ones and degrade them by lysosomal enzymes (Overholtzer et al., 2007). This process can occur when cancer cells struggle to scavenge nutrients from their microenvironment, particularly where tumor vasculature is either deficient or absent – like in organoids. Interestingly, entosis has yet to be described in colorectal cancer (Fais and Overholtzer, 2018), and unlike previously reported (Overholtzer et al., 2007), exposure to Matrigel did not inhibit the internalization process.

To provide comprehensive characterization of the heterogeneity at the subcellular level across the different tissue morphologies, we proceeded to annotate cells and organelles by segmentation of the FIB-SEM data. While manual annotation becomes unfeasible given the large amount of data, deep-learning algorithms offer a workable solution for automating this task. We used convolutional neural networks (CNN) provided in the software ORS Dragonfly and trained instance-type segmentations networks for single cellular structures (Material and Methods). We focused on two commonly segmented targets (cell nuclei and mitochondria) that are large, have a roughly defined shape and are highly represented in the cell volumes, and two challenging fine structures (actin bundles in microvilli and cell junctions). Manual annotation was first performed on multiple subframes to train a CNN that can automatically segment a specific organelle (e.g., mitochondria) within the whole volume (Fig 6A-C). Of special relevance for cancer development are the interactions between cortical cytoskeleton and cell junctions (te Boekhorst and Friedl, 2016). Junctional complexes play a role in the oncogenic signalling cascade which regulates cytoskeleton remodelling. This leads to a loss of epithelial cell polarity and tissue invasion by cancer cells in the process of epithelial-to-mesenchymal transition (Huang et al., 2022). Despite the low amount of stain (0.1% uranyl acetate) in our preparations, patient-derived colorectal cancer organoids featured high contrast for actin bundles and different types of cell junctions. Actin bundles were mostly evident within microvilli at the apical side of cells facing a lumen, appearing as ∼1µm long fibers or high contrast dots when cut parallel or perpendicular to the image plane, respectively (Fig 6D, Movies 2-4). The trained CNN segmented scattered actin bundles within cells and those that form the basis of luminal microvilli (Fig 6E-F and Movie 5). Cell junctions are inherently smaller and of lower representation in the data. Although the FIB-SEM resolution was sufficient to distinguish between tight junction “membrane kissing points” (Fig 6G, H white arrows) and larger desmosomes (Fig 6H, red arrows), the CNN could not segment them separately due to their insufficient pixel sampling. Nonetheless, a dedicated CNN segmented the detectable cell junctions in all the EM data (Fig 6J, K). A 3D rendering of the cell nuclei, actin bundles and cell junctions of one of the acquired volumes (acquisition n°3 in Fig S6A) shows the complexity of the organelle arrangement in colorectal cancer organoids (Fig 6L).

**Figure 6.**
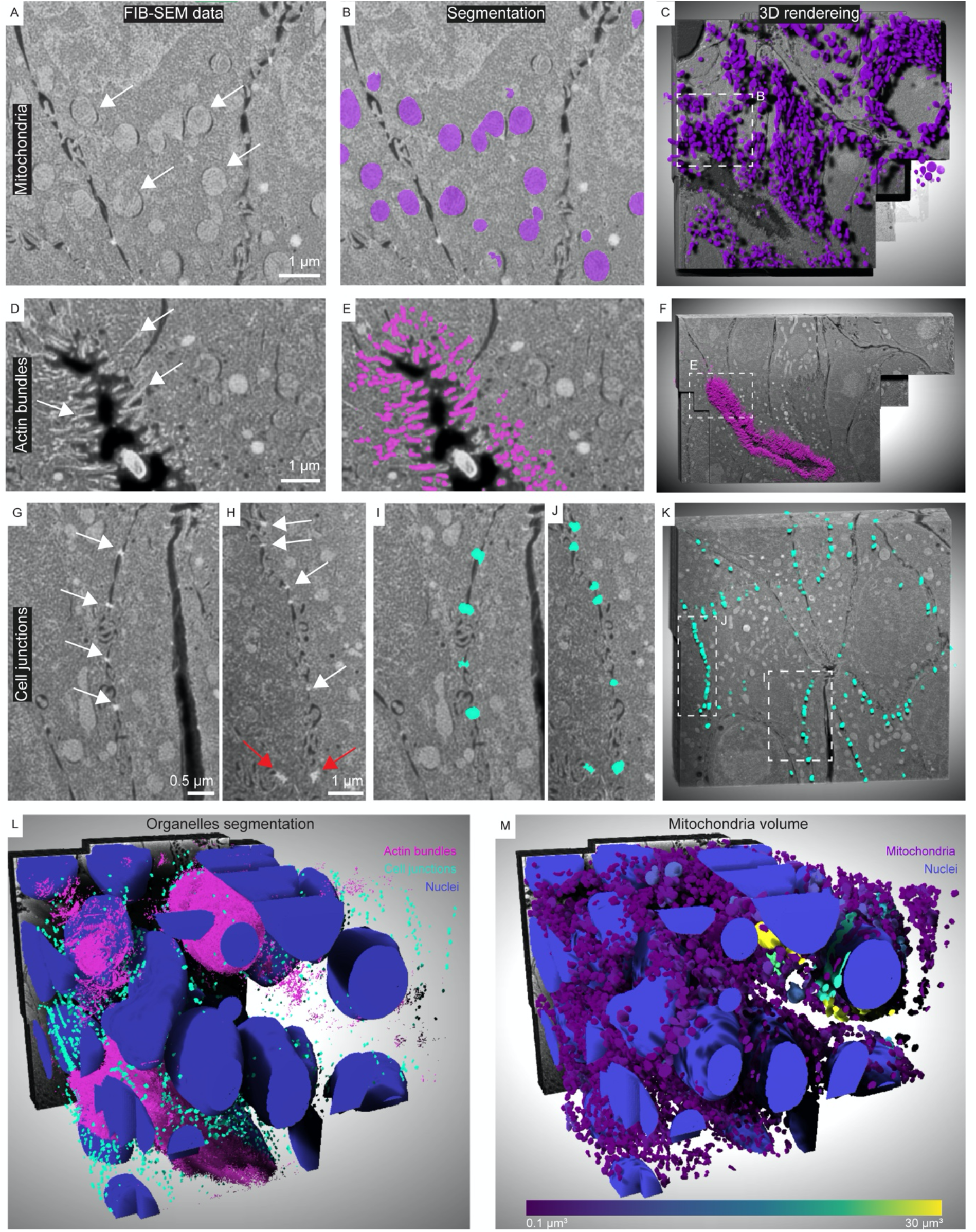
FIB-SEM volumes of patient-derived colorectal cancer organoids reveal fine ultrastructural details. **A**, FIB-SEM frame showing mitochondria (white arrows), overlayed (**B**) with the corresponding automatic segmentation. **C**, 3D rendering of 50 consecutive frames with embossed mitochondria segmentation (purple). Frame indicates the area shown in **A, B. D**, details of actin bundles in microvilli (white arrows) from a single frame of a FIB-SEM acquisition and the corresponding automatic segmentation (magenta, **E**). **F**, 3D rendering of 50 consecutive frames with embossed actin bundles segmentation (pink). Frame indicates the area shown in **D, E. G, H**, details of cell junctions (white arrows) and desmosomes (red arrows) from a single frame of a FIB-SEM acquisition and the corresponding automatic segmentation (**I, J**). **K**, 3D rendering of 50 consecutive frames with embossed cell junctions segmentation (cyan). Frame indicates the areas shown in **G-J. L**, 3D rendering of automatic segmentation of subcellular structures: nuclei (blue), actin bundles (magenta) and cell junctions (cyan). Volume corresponds to the dataset in position 1, Figure S6A. **M**, 3D rendering of automatic segmentation with nuclei (blue) and mitochondria colored according to their volumes (purple to yellow gradient); note the two-order of magnitude larger mitochondria around the nuclei of two cells (green-yellow).

The segmentations of the different structures make it possible to perform statistical analysis of the organelles for quantitative characterization of the organoids. For example, mitochondria generally range from onion-like to highly reticular shapes, whereas their dimensions are roughly uniform in non-diseased cells (Egner et al., 2002; Fawcett, 1966; Schmidt et al., 2009). When scoring mitochondria according to their volume, we found that they overall showed consistent volumes with the exception of one of the volumes (n°3 in Fig S6A) featuring two cells with mitochondria that were two orders of magnitude larger than the others (Fig 6M). However, we could not identify structural features in the cells or their environment that otherwise point to their different state.

We next turned to quantitatively evaluate differences in subcellular structures associated with the different cell packing arrangements captured in our three FIB-SEM acquisitions. Based on the fluorescence data, we refer to the different cell packing as mixed, monolayer or compact (Fig 7). The mixed area shows heterogeneous arrangements (Fig 7A, green; acquisition n°3 in Fig S6A); a crypt displays a long lumen surrounded by cell monolayers (Fig 7A, red; acquisition n°2 in Fig S6A); the compact zone shows dense cell packing (Fig 7A, blue; acquisition n°1 in Fig S6A). It is known that healthy polarized cells in epithelia tend to have roughly spherical nuclei and asymmetric organelle distribution in 2D cultures (Skinner and Johnson, 2017). It is however expected that cancer cells will exhibit more heterogenous morphologies that have yet to be described in high detail in 3D cell cultures. Indeed, the colorectal cancer cells show markedly spherical nuclei in mixed cell-packing parts of the organoids (Fig. 7B, green), while monolayer or compact cell-packing reveal spheroid-shaped nuclei (Fig 7B, red and blue) with equal distribution between oblate and prolate orientations with respect to the imaging direction. This is in line with literature from 2D cell cultures where cancer cells adapt nuclear shape to account for external forces or in response to constrained space during invasion (Zink et al., 2004). Interestingly, the ratios between nuclei and cell volumes, and mitochondria and cell volumes were similar in all three cell-packing morphologies (Fig 7H). However, the majority of cells in our acquisitions show nuclei with long clefts and grooves, and occasionally polynucleated cells with mitochondria intertwined between the nuclei (Movies 5-7), demonstrating the heterogenous nature of the organoids that can only be pertained in 3D conditions. We then examined whether a relationship existed between multicellular architectures and the spatial distribution of cell junctions (Fig. 7C-F). We counted 7.803, 4.980 and 5.105 individual cell junctions in the areas with mixed, monolayer and dense cell packing, respectively (Fig 7H). A visual inspection of the segmentations showed that the cell junctions nicely follow the cell boundaries (Fig 6K and Movies 5-7). Nearest-neighbour analysis of the segmented cell junctions interestingly showed that inter-junction shortest distances in all three volumes have a similar pattern, with 90% of the data falling within 1 µm distance (Fig 7C). However, global nearest neighbour analysis does not account for local variations when evaluating the entirety of the multicellular volume. We therefore calculated density maps of the local concentration of cell junctions (Materials and Methods) (Fig 7E) and for the actin bundles (Fig 7F). The density maps indicate that local higher density of cell junctions correlates with local higher density of actin bundles that line up the organoid lumina. Our data demonstrates that more compact and polarized epithelia seal off lumina with a higher local concentration of cell junctions and microvilli, reminiscent of patterns observed in the MDCK (Madin-Darby canine kidney) cell-line model using light microscopy and thin sectioning TEM (Beutel et al., 2019; Odenwald et al., 2018; Otani et al., 2019). While the molecular composition of junctional complexes and their dynamicity are well characterized in homogenous cell-line models that can establish epithelial polarity, not much is known about their 3D spatial arrangement at ultrastructural resolution in heterogenous organoids. Our data and analyses thus provide the first quantitative description of the local distribution of diffraction-limited cell junctions in cancer tissues models using patient-derived organoid cell cultures.

**Figure 7.**
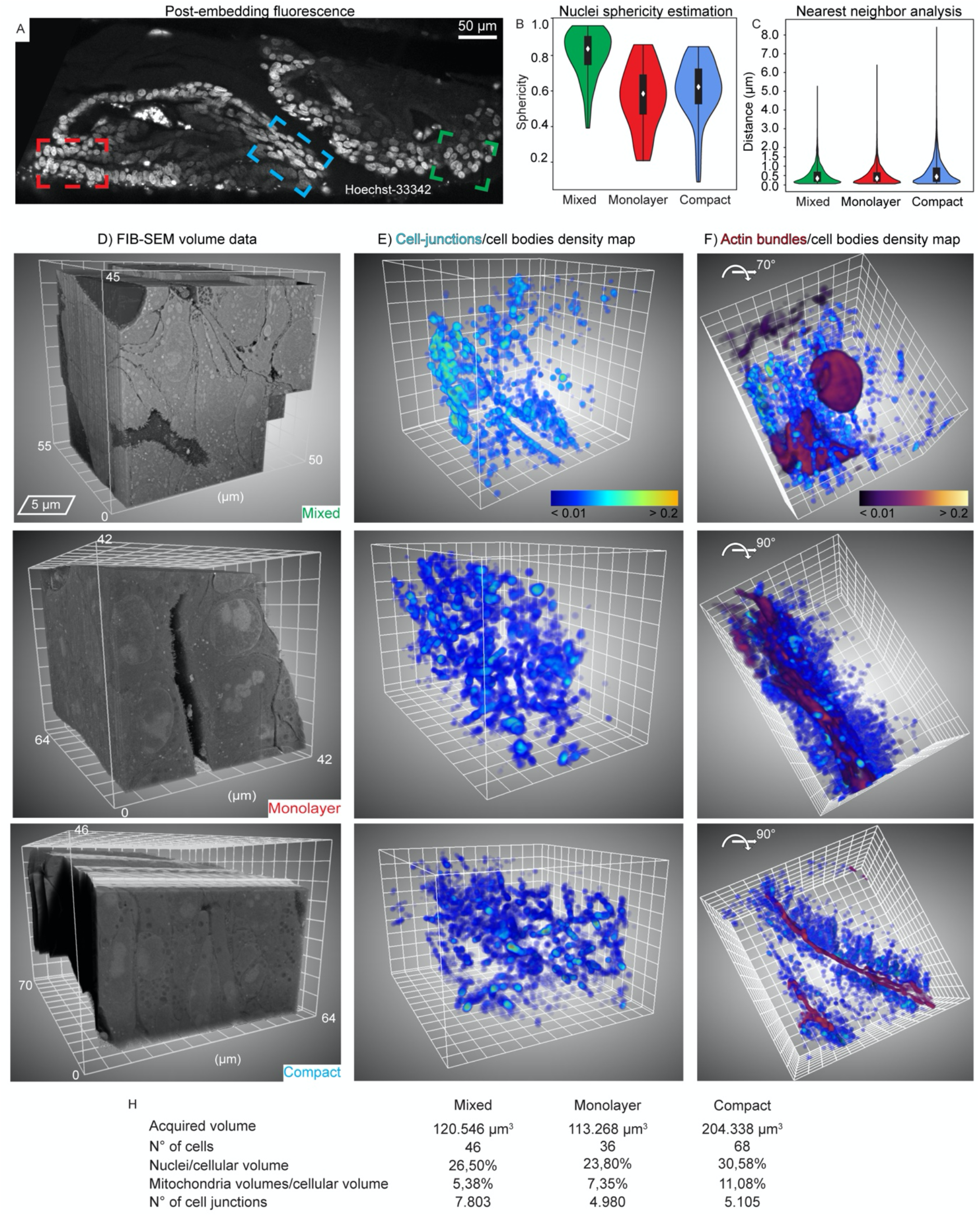
Quantitative analysis of fine cellular morphology in patient-derived colorectal cancer organoid. **A**, a confocal plane from patient-derived colorectal cancer organoids (volume rendered in Figure S6C). Post-embedding fluorescence shows cell nuclei stained with Hoechst (white). Highlighted are the FIB-SEM acquisitions corresponding to different cell-packing morphologies: mixed (green), monolayer (red) and compact (blue) corresponding to positions 3, 1, 2, respectively in Fig. S6A. **C**, nuclear sphericity and **B**, nearest neighbour analysis of cell junctions based on the automatic segmentations exemplified in Figure 6. **D**, 3D representation of the three morphologically-different areas. x,y and z dimensions are shown. Detailed overview of the FIB-SEM volumes and corresponding organelle segmentations are shown in Movies 2-7. The first 20 µm of the mixed morphology (**D**) are not displayed to aid the visualization of multiple lumina. **E**, local density maps of cell junctions based on the automatic segmentation colored as a blue-orange gradient (mesh size 5 µm). **F**, same as **E**, rotated by 90 degrees, and include the actin bundles density maps (fire gradient). **H**, summary of the salient statistics of the acquired FIB-SEM volumes.

## Discussion

We describe a multiscale imaging pipeline across optical and electron microscopes that enables seamless correlative investigations in the emerging model systems of 3D cell culture. Performing the 3D culture directly in standard HPF carriers presented several advantages, and at the same time retained the expected morphology for a diverse set of model organoids (Fig 2). First, it avoids manual transfer of organoid samples embedded in a fragile gel from standard TC culture dishes to the specialized carriers for cryo-fixation. This guarantees that the sample remains unperturbed from the beginning of cell seeding until high-pressure freezing. Second, it avoids the need for dedicated carriers (Hötte et al., 2019) or additional medium embedding for mounting for confocal and light-sheet applications (Huang et al., 2021). Therefore, the use of HPF carriers enabled us to correlate the growth of the 3D cell culture (stereo-microscopy), the visualization of multicellular architectures and cellular structures of interest (live-cell fluorescence confocal imaging) for subsequent post-embedding imaging and two-photon laser branding, to ultimately guide FIB-SEM volume imaging for ultrastructural investigations. As proof-of-principle, we also demonstrated that it is possible to perform fluorescence light microscopy in cryogenic regime on HPF carriers (Fig 3H, I). Cryo-CLEM for HPF samples still suffers from technical shortcomings, including light scattering when imaging beyond the first few microns from the HPF carrier surface, thus requiring new objectives with working distance ≥ 1mm and with NA > 0.5. In addition, unlike room-temperature approaches, it is not possible to brand the sample surface for precise correlation with FIB-SEM imaging, although new carriers with finder patterns may provide a possible solution (Kapteijn et al., 2022).

HPF carriers further provide a high surface area/volume ratio which allows infiltration of the live cultures with diverse small molecules (100 to 1000 Da molecular weight), including drugs like doxycycline (Fig S1) and commonly used live dyes (Fig S2). Thus, 3D cell cultures performed in HPF carriers are amenable to drug screening and treatment, with the advantage of being directly accessible to imaging by both light and electron microscopy. Because of the unspecific and broad-spectrum binding of the uranyl-acetate stain, FIB-SEM can image every sub-cellular component at nanometer-scale resolution within hundreds of thousands of cubic microns of a multicellular sample guided by fluorescent signal. Future efforts for integrating such approaches into high-throughput platforms being developed for organoid research will facilitate expanding organoid phenotyping to the ultrastructural level and aid in deriving mechanistic understanding of tissue-specific processes or drug effects.

We establish that several live dyes are resilient to freeze-substitution. Rochi et al., (Ronchi et al., 2021) recently demonstrated that it is possible to exploit the fluorescent signal of transgenic fusion proteins to target FIB-SEM acquisitions. However, the introduction of transgenic fluorescent reporters is time consuming and, may be impractical in case of patient-derived organoids aimed at speedy translational molecular medicine. Therefore, small molecule live dyes that bind specifically to cellular structures can be used to identify specific cells or phenotypes (Day and Davidson, 2009), paving the way to study any culture system that is not easily accessible to genetic engineering. It is important to highlight the limits of light microscopy for large and opaque objects like organoids. Photon scattering within biological material increases with the light path through the sample. Thus, fluorescence signal becomes dimmer the deeper one needs to image (Helmchen and Denk, 2005; Ntziachristos, 2010). This problem might be enhanced by limited diffusion of cell-permeable fluorescent dyes within a compact, tight-junction sealed epithelium. As a result, loss of fluorescence signal intensity is particularly severe for structures with dense cell packing like spheroids and parts of colon organoids (Fei et al., 2022; Richter et al., 2022). We observed that the fluorescence signal exhibits a gradient from the core of such organoids e.g., Fig 3C and Fig 7A.

Optimizing the freezing condition is necessary for each HPF sample type (Möbius et al., 2010). We used broad tiled TEM acquisitions of 2D sections as a straightforward analytical tool to survey the different organoid samples for ultrastructural preservation and to optimize freezing conditions. ECM-embedded 3D cell cultures are difficult to vitrify, especially in the case of cystic organoids with large lumina. In our hands, mouse mammary gland organoids showed the most serious freezing artifacts. We show that it is possible to improve HPF cryo-fixation for this system by using anti-freeze additives, such as the ready-to-use Cellbanker 2 culture medium and 20% Ficoll diluted in standard cell culture medium, establishing a useful reference for future studies on cystic organoids.

The ability to image 3D cell cultures across scales allowed us to investigate diffraction-limited structures in their multicellular context that are otherwise difficult to capture by either 2D TEM or light microscopy alone. Fluorescence signal guided the FIB-SEM acquisition of more than 10^5^ cubic microns of the colorectal cancer organoid volume at ultrastructural resolution, corresponding to roughly 30 to 70 cells in a single imaging session. Further, automatic image segmentation provided high-throughput and quantitative information on the imaged data. The combination of these methods provides the first ultrastructural 3D map of cell junctions using organoid models. The data unveiled that cancer cells sealing off lumina, despite exhibiting some features of healthy cells (like microvilli), displayed abnormal polarization. This could result from some cells with higher invasive and metastatic potential within different parts of the tumor. Such level of tumor heterogeneity can only be probed at ultrastructural resolution, across multiple cells within an epithelium.

Beyond the scope of this study, potential applications of our pipelines are several and multidisciplinary. We envision that patient-derived tumor organoids will be amenable to ultrastructural analysis to study multicellular interaction during drug treatment, enabling a deeper understanding of drug resistance e.g. via loss of polarity/epithelial-mesenchymal transition (Croix and Kerbel, 1997; Dongre and Weinberg, 2019; van Staalduinen et al., 2018) and therapies modulating cell-cell contacts (Vucetic et al., 2020). Co-cultures of patient-derived colorectal carcinoma organoids have been further used to interrogate the effects of anaerobic microbiota on tumor progression (Pleguezuelos-Manzano et al., 2020). The use of fluorescently marked patient-derived organoids together with differentially marked associated microbial taxa allows now the ultrastructural investigation of detrimental tissue/bacteria interactions (Schmidt et al., 2018) as well as the detrimental loss of epithelial barrier integrity (Yu, 2018). Furthermore, we demonstrated the possibility to assess mitochondria shape and volume, which can foster research on mitochondrial fusion and fission (Adebayo et al., 2021) and mitochondrial abnormalities in the context of a 3D organoid organisation. The mitochondrial form and resulting (dys)function play an important role in tissue homeostasis and disease, including in several solid cancers (Nunnari and Suomalainen, 2012), which can be assessed with the appropriate 3D organoid models.

In summary, the use of 3D culture technology to model tissue complexity and cellular crosstalk is becoming increasingly evident, but lacks appropriate multimodal tools for comprehensive studies across spatial scales. Here we developed a multiscale imagine pipeline achieving a resolution continuum from millimeter to nanometer scale, and show its applicability to four common 3D cell cultures used in organoid and cancer research. While sample-specific optimization will be required to achieve optimal ultrastructural preservation for different organoid types, the developed pipeline and protocols could be easily adopted by EM labs that routinely practice HPF and FIB-SEM volume imaging, broadening their capability to the study of 3D cultures. The combination with live-cell dyes permits the selection of regions of interest that can be followed in ultrastructural resolution. We envision that our pipeline further establishes the basis for applications in full cryogenic regime with the aim to expand the resolution range to the molecular scale.

## Materials and Methods

### Animal experimentation

Animals are treated at the European Molecular Laboratory in agreement with National and International laws and policies. All efforts are made to use the minimal possible amount of animals in accordance with Russell and Burch’s (1959) principle of (3Rs) reduction and highest ethical standards. The work was approved by the IACUC (Institutional Animal Care and Use Committee, approval #160070 to MJ).

### Animals

TetO-MYC/ MMTV-rtTA (D’Cruz et al., 2001) and TetO-Neu/ MMTV-rtTA (Moody et al., 2002) mouse strains were bred to obtain TetO-MYC/ TetO-Neu/ MMTV-rtTA animals. A reporter R26-H2B-mCherry mouse strain line (Abe et al., 2011) (RIKEN, CDB0239K) was crossed in to establish the experimental line TetO-MYC/ TetO-Neu/ MMTV-rtTA/ R26-H2B-mCherry in FVB background. The housing of all animals used in this study was performed in the Laboratory Animal Resources facility at EMBL Heidelberg, according to the guidelines and standards of the Federation of European Laboratory Animal Science Association (FELASA).

### 3D Cultures

All sample typs described hereafter share culture conditions in both HPF carriers and standard TC dishes, as well as handling, only differing in the Matrigel volume and the carrier type.

### Primary Mouse Mammary Epithelial Cells

Mammary glands were dissected from 7-9 weeks old female virgin mice and collected in a 15 ml falcon. For dissociation of the tissue, the mammary glands were digested overnight at 37 ºC and 5% CO_2_ in a loosely capped 50 ml falcon with 5 ml of DMEM/F12 (Lonza) supplemented with 25 mM HEPES, 1% Penicillin Streptomycin solution (ThermoFisher), 750 units of Collagenase Type III (Worthington Biochemical Corporation) and 20 µg of Liberase TM (Roche). After 16 hours digestion, cells were washed with DMEM/F12 (Lonza) supplemented with 25 mM HEPES, 1% Penicillin Streptomycin solution (ThermoFisher). Pellet was then trypsinized with 5 ml of 0.25% Trypsin-EDTA (ThermoFisher) to obtain single cells following the published protocol (Jechlinger et al., 2009). To generate 3D cultures, a master mix of Matrigel Growth Factors Reduced (Corning, 356231), Rat Collagen I (RnDSystems, 3447-020-01), and PBS was prepared in ice following 4:1:1 proportion. The cell suspension and matrix mix were combined to obtain a concentration of 30,000 cells/100 µl of master mix. Volumes of 1 µl were seeded in the 200 µm deep HPF carriers, previously sterilised with 70% Ethanol and fixed with Matrigel (Corning) to an 18 mm glass coverslip. Then, each carrier was placed into a well of a 24-well plate (Corning) to allow the gels to polymerize at 37 ºC with 5% CO_2_. After polymerization, each gel was supplied with 1 ml of Mammary Epithelial Cell Growth Medium (Promocell, c-21010) supplemented with Mammary Epithelial Cell Growth supplement (Sciencell, 7652) and incubated at 37 ºC in a humidified atmosphere with 5% CO_2_. For induction of oncogene expression in the organoids, Doxycycline Hyclate (Sigma, D9891) was diluted into the growth medium at 600 ng/ml after 4 days of 3D cell culture. Growth media were exchanged every other day in both conditions.

### Human Organoid Culture

Biosamples were provided by Lungbiobank Heidelberg member of the biomaterial bank Heidelberg (BMBH) in accordance with the regulations of the BMBH and the approval of the ethics committee of the University of Heidelberg (study S-270/2001 -biobank vote). Metastatic lesion from a patient diagnosed with metastatic colorectal cancer was surgically removed and dissected from lung tissue. A tumor fragment of a minimum of 100 mm^2^ was used for the establishment of primary organoid culture. Briefly, tissue was mechanically dissociated using scalpel followed by pipetting through a 10 ml pipette in a basal medium Advanced DMEM/F12 (Gibco), Primocin 50 µg/ml (Invivogen), 1% GlutaMAX (Gibco), 1% HEPES (Gibco), 1% Penicillin Streptomycin solution (Gibco), N-Acetylcysteine 1.25 mM (Sigma) supplemented with 1% B27 supplement (Gibco). Next, the cell suspension was enzymatically digested for 2 hours at 37 ºC in basal medium supplemented with Liberase DH (final concentration 0.28 Wünsch units/ml). The cell suspension was then filtered through 100 µm and 40 µm cell strainer. Single cells were seeded in Growth-Factor Reduced Matrigel (Corning) mixed with PBS (4:1) adjusted to a concentration of 17,000 cells/µl and cultured in culture medium Advanced DMEM/F12 (Gibco), Primocin 50 µg/ml (Invivogen), 1% GlutaMAX (Gibco), 1% HEPES (Gibco), penicillin 100 U/ml and streptomycin 100 μg/ml (Gibco), N-Acetylcysteine 1.25 mM (Sigma) supplemented with 1% B27 supplement (Gibco), 50 ng/ml Epidermal Growth Factor (EGF) (Peprotech), 100 ng/ml Noggin (Peprotech) and 500 nM A83-01 (Tocris Bioscience). Organoids were passaged with Gentle Cell Dissociation Reagent (StemCell Technologies) according to manufacturer instructions. For imaging experiments, organoids from passage 4-10 were seeded in HPF carriers using 2 µl of organoid suspension in Matrigel/PBS (6:1).

### BT474 cell spheroids

BT474 human cell line was obtained from American Type Culture Collection. Cells were grown in DMEM 1x 4.5 g/L D-glucose FluoroBrite (Gibco) medium supplemented with 10% inactivated FBS (Gibco), 1% HEPES 1 M (Gibco), 1% Sodium Pyruvate 100 mM (Gibco), 1% MEM NEAA 100X (Gibco), 1% L-Glutamine 200 mM (Gibco), 1% Penicillin Streptomycin solution (Gibco) and passaged using 0,05% Trypsin-EDTA (Gibco). For spheroid formation and imaging experiments, cells were seeded in HPF carriers in Growth-Factor Reduced Matrigel (Corning) mixed with PBS (6:1) at 500 cells/µl.

### Immunofluorescent staining

Organoid cell culture was performed in HPF carriers (Wohlwend GmbH) which were then transferred to a deactivated clear glass vial (Waters,186000989DV), and fixed with 4% PFA for 5 minutes followed by three washes of PBS. To prevent nonspecific antibody binding, the HPF carriers were incubated with 10% goat serum for 2 hours at room temperature. Primary antibody incubation was performed overnight at 4 ºC. Afterward, HPF carriers were washed with PBS three times for 10 minutes, and incubated with secondary antibodies and DAPI (Thermo Fisher, 62248, 1:1000). Samples were mounted using Prolong Gold with DAPI (Thermo Fisher, P36931). The following antibodies were used in this study: c-MYC (Cell Signaling Technologies, 5605, 1:800), ZO-1 (Thermo Fisher, 61-7300, 1:400), E-cadherin (Thermo Fisher, 13-1700, 1:200), Alpha6-integrin (Millipore, MAB1378, 1:100), Cytokeratin 8 (Troma-I, DSHB, 1:100), Cytokeratin 14 (Invitrogen, MA5-11599, 1:200) Cytokeratin 20 (Abcam, ab76126, 1:200), Alexa 488 (Invitrogen, A11034, 1:800), Alexa 568 (Invitrogen, A11031, 1:800), Alexa 647 (Invitrogen, 21247, 1:800). Mounts were imaged in HPF carriers on a Leica SP5 confocal microscope using a 63x 1.2 NA water immersion lens and the LAS AF imaging software. HPF carriers were mounted facing the objective on top of a Nunc Lab-Tek II chambered 1.5 borosilicate coverglass.

### Stereomicroscopy and widefield transmission imaging

The 3D cell cultures in HPF carriers were maintained in 24-well plates and imaged in a Leica M125 C Stereomicroscope in dark field mode to enhance the HPF carriers contrast against the background. 3D cell culture grown in gels in TC-dishes over the time-course of the experiment were imaged using the widefield high-throughput Olympus ScanR microscope in transmission mode. Each well of the 24-well plate was imaged using 9 ROI per well, with 21 Z-stacks (100 µm of scanning step in Z). Images were acquired with 4x UplanSApo 0.16 NA Air objective in an environmental chamber at standard cell culture conditions (37 °C, 5% CO_2_). Projections of z-stacks and image stitching were done using Fiji (Schneider et al., 2012).

### Live-cell confocal imaging

Here, we refer to single time-points acquisitions and not to time-series acquisitions commonly known as 4D imaging. For our live cell imaging, cultures in HPF carriers were washed with PBS once, transferred to a 35 mm or 10 mm diameter cell culture dish (Greiner Bio One International, catalog number 627860, Kremsmünster, Austria). For samples devoid of genetic fluorescent tags, the HPF carriers were incubated for 20 minutes with the desired live dyes diluted in the growth medium to the following final concentrations: SiR-actin 100 µM (Spyrochrome AG, catalog number SC001, Stein am Rhein-Switzerland), Hoechst-33342 10 µM (Thermo Fisher Scientific, Waltham, MA USA, catalog number H1399), FM4-64 2 µM (Thermo Fisher Scientific, Waltham, MA USA, catalog number T13320) and BODIPY 493/503 1 µM (Thermo Fisher Scientific, Waltham, MA, catalog number D3922). The culture dish was connected to the microscope objective by a drop of deionized water, and HPF carriers were flipped with the recess facing the microscope objective lens. Live cell imaging was performed at 37°C and 5% CO_2_ with a Zeiss LSM 780 NLO (Jena, Germany) equipped with a LD LCI Plan-Apochromat 25x 0.8 NA water immersion objective. To image the whole HPF carrier volume and locate single organoids, we first detected cell nuclei marked with Hoechst-33342 and acquired z-stacks with 6×6 tiles of 512^2^ pixels with 10% overlap, and 10 μm z-step. The final montage was stitched using ZEN-black software. For single organoids, we acquired z-stacks of 1024^2^ pixels at different z-steps ranging from 1.0 or 1.8 μm.

### High-pressure freezing, freeze-substitution, and two-photon laser branding

We followed the procedure described in (Ronchi et al., 2021). Briefly, we high pressure froze 3D cell cultures in HPF carriers with an HPM 010 (AbraFluid). When necessary, before high-pressure freezing the carriers containing the 3D cell cultures were dipped for one minute either directly in Cellbanker 2 cryo-preserving medium (catalog number 11891, AMSBIO, Cambridge -US) or in 20% Ficoll 70.000 MW (catalog number F2878, Sigma, Merck KGaA, Darmstadt, Germany) diluted in Mammary Epithelial Cell Growth Medium (Promocell, c-21010). Next, we performed freeze-substitution with 0.1% uranyl acetate (UAc) in acetone. After 72 h incubation at -90 °C, the temperature was increased to allow the reaction of UAc with the biological material. We then rinsed the samples with pure acetone before infiltration of the resin lowicryl HM20 (Polysciences). The resin was polymerized with UV at -25 °C. The resin-embedded samples were then transferred in 35 or 10 mm cell culture dish (Greiner Bio One International 627860, Kremsmünster, Austria) with water as immersion medium and imaged at an inverted Zeiss LSM 780 NLO microscope (Jena, Germany) equipped with a 25x Plan-Apochromat 25x 0.8 NA Imm Korr DIC multi immersion objective lens. Surface branding was performed with the 2-photon Coherent Chameleon Ultra II Laser (Santa Clara, USA) of the Zeiss LSM 780 NLO microscope and the “bleaching” function of ZEN black software.

### Cryo-Confocal microscopy

Confocal stack acquisition under cryo-conditions was carried out with Leica TCS SP8 upright microscope (Leica microsystems CMS GmbH, Mannheim, Germany), controlled by Leica Application Suite X 3.5.5.19976 software. The microscope was equipped with cryo-stage, insulated HC PL APO 50x 0.90 NA DRY and HC PL FLUOTAR 50x 0.80 NA objectives, and Leica DFC365 FX camera. The high-pressure frozen organoids in HPF carriers were inserted into the shuttle (Leica microsystems CMS GmbH, Mannheim, Germany) with the sample side facing the objective. All sample loading and transfer operations were conducted in the liquid nitrogen vapor phase using a dedicated loading/transfer unit (Leica microsystems CMS GmbH, Mannheim, Germany). The loading and transfer steps were carried out in a humidity-controlled room (10%) to minimize ice contamination on the sample. After the sample was deposited on the microscope stage pre-cooled to -195° C, we determined the approximate z-position of the sample surface in wide-field illumination mode. Using Matrix MAPS+CLEM2 application within the Matrix Screener module, we acquired 100 μm deep (steps of 2 μm) z-stack tiles (x/y with 20% overlapping) to produce a stitched 3D overview image of the whole carrier at the desired wavelengths (Leica Application Suite X 3.5.5.19976). This helped to determine ROIs for subsequent confocal stacks acquisition. Confocal stacks were acquired in two channels: 405 nm at 50% power Diode laser (HyD1 detector 410 nm - 504 nm) and 552 nm at 50% power OP SL laser (HyD 3 detector 660 nm - 778 nm). Pinhole 1 AU, zoom 1, pixel format 1056×1056, pixel size in xy = 0.22 μm, in z = 0.372 μm, bidirectional scan at 200 Hz. Z-stack of 50 μm depth were acquired with a z-step system optimized for the 405 nm wavelength. Confocal stacks were processed with the Lightning deconvolution software module (Leica Application Suite X 3.5.5.19976) at default settings and a number of iterations set to 5 for both channels. After the acquisition, the HPF carriers were retrieved under cryo-conditions for further processing.

### FIB-SEM

After confocal imaging and branding for targeting, the blocks were mounted on a SEM stub using silver conductive epoxy resin (Ted Pella). After mounting, the blocks were gold sputtered with a Quorum Q150R S coater. FIB/SEM imaging was performed on a Zeiss CrossBeam XB540 or XB550. Briefly, platinum was deposited over the area marked by the laser branding. Auto-tuning marks were milled on the platinum surface and highlighted with carbon. Large trenches were milled with 30 nA FIB current and surface polished with 7 nA or 15 nA. Precise milling during the run was achieved with currents of either 700 pA or 1.5 nA. For all experiments, the imaging was done with an acceleration voltage of 1.5 kV and a current of 700 pA, using a back-scattered electron detector. For single cells within organoids, data were acquired at 10 nm isotropic (x,y,z) sampling. For the acquisition of entire organoids, voxels of 15×15×20 nm^3^ were found to be the best compromise between the achievable resolution/field-of-view ratio and milling stability.

### Thin sectioning TEM

Blocks prepared as described above were sectioned with an ultramicrotome (Leica UC7) and 70 nm sections were collected on formvar coated slot grids. Organoid TEM images were acquired without post-staining using a Jeol 2100 Plus operated at 120 kV. The organoids were identified at 400x mag (32 nm pixel size). Subsequently, we imaged single organoids by fitting a montage of tiles at 2000x mag (6.7 nm pixel size) using SerialEM (Schorb et al., 2019). The montages were stitched with the command justblend and, where necessary, manually corrected with Midas, both implemented in Imod (Kremer et al., 1996).

### Deep learning automatic volume segmentation and data-mining

ORS Dragonfly 2021.3 (Object Research Systems, Quebec, Canada) and subsequent versions were used to train neural networks for automatic segmentation of FIB-SEM data of organoids subcellular structures. Dragonfly was installed on a virtual machine workstation with Windows 10 pro mounting an Intel(R) Xeon(R) Gold 6226R CPU 2.90GHz with 128 GB RAM and an Nvidia V100S card with 30 GB RAM. For pre-processing, the raw FIB-SEM frames were first aligned with the Scale-Invariant Feature Transform (SIFT) algorithm (Lowe, 1999) implemented as a macro in Fiji (https://imagej.net/plugins/linear-stack-alignment-with-sift) (Schneider et al., 2012) allowing only image transformation by translation. Next, the images were cropped to exclude pixels without signal outside imaged areas. FIB-SEM data sets containing few cells acquired with 10 nm isotropic resolution were processed as such, while whole organoid acquisitions (15×15×20 nm resolution) were binned twice for computational efficiency. CNN models were prepared with the 2.5D-UNet architecture and the following design: depth level 6 with 7 slices for CNNs for actin bundles, nuclei, cell bodies and cell boundaries; depth level 6 with 5 slices for cell junctions and depth level 5 with 5 slices for mitochondria. Next, using the segmentation wizard module, ground truth data was manually annotated from 2 to 5 subframes of each dataset and used to train each CNN with 10-fold data augmentation by: flipping vertically and horizontally, rotation up to 180°, shear up to 2° and scaling between 90 and 110%. For training, patches ranging between 128^2^ and 512^2^ pixels were used depending on the size of the objects. These initially trained CNNs models were then used to segment 5 to 10 whole frames from each dataset, where errors were corrected manually. Then, the model weights were reset and the CNNs were re-trained using the whole selected frames without data augmentation this time. Finally, the CNNs were run over all the datasets to segment cell bodies and subcellular structures. To compute the nearest-neighbour analysis, the instance segmentation of the organelle of interest was first converted to “multi-ROI”. Here, the global instance segmentation is split into single 3D objects based on their 3D connected components: objects not connected to each other are considered as an independent item belonging to a global instance segmentation. Due to the lower resolution in z of the FIB-SEM images used for segmentation (40 nm), objects closer than 50nm were excluded by the count. Then, the nearest-neighbour was calculated using the “Find minimum distance between objects” option. To compute local density maps of segmentations, “Bone analysis” wizard was used to estimate volume fractions. After optimizing parameters for resolving power and computing time, each FIB-SEM acquisition was sampled using a sphere of 1 µm diameter (∼ 4 µm^3^ sampling volume) and scanning the volume using 0.25 µm step size. The resulting density maps were colored and displayed in Dragonfly. All the statistical analysis was plotted using Matplotlib (Hunter, 2007).

### Data processing and rendering

Unless otherwise specified, all microscopy data was processed with Fiji (Schneider et al., 2012). Where necessary, the contrast of electron microscopy data was enhanced using the Fiji plugin Enhance Local Contrast (CLAHE) using the following parameters: blocksize 63, 256 histogram bins and maximum slope of 2.0. When necessary, before contrast enhancement, FIB-SEM volumes were corrected for curtaining effect applying wavelet decomposition (Münch et al., 2009) implemented in the open-source software SerialFIB (Klumpe et al., 2021).

Here, coif3 type vertical wavelets with a decomposition sigma level of 8 and a sigma gaussian for vertical stripes dampening of 6 were used. All FIB-SEM and segmentation rendering and movie making was done with ChimeraX (Pettersen et al., 2021). For light microscopy data 3D rendering, Imaris 9.6.0 with either “blend” or “shadow projection” rendering modes was used. Movies with multiple channels synchronized were made with the Fiji macro by Patrice Malscalchi (https://github.com/AiviaCommunity/ImageJ-Macros-Utilities). Schematics were created with Biorender (Biorender.com) and Adobe illustrator.

## Acknowledgments

We acknowledge support from the EMBL Electron Microscopy Core Facility, Advanced Light Microscopy facility, IT Services, especially Thomas Hoffmann, and the EMBL Laboratory for Animal Resources. We thank Davide Floris for discussions about image processing, Martina Dees for kindly providing Hoechst-33342, Mark Schneider and Christa Stolp from Thoraxklinik University Hospital Heidelberg for providing tissue samples for deriving human colorectal cancer organoids. The prototype cryo-confocal fluorescence light microscope was developed in collaboration with Leica Microsystems. EDI was supported by a fellowship from the EMBL Interdisciplinary (EI4POD) programme under Marie Skłodowska-Curie Actions COFUND (847543). SG was supported by a fellowship from the EMBL Interdisciplinary (EI3POD) programme under Marie Skłodowska-Curie Actions COFUND (664726). JM acknowledges funding from the EMBL and the European Research Council (ERC 3DCellPhase^-^ 760067).

## Authors contributions

EDI: Conceptualization, methodology, Validation, Formal analysis, Investigation, Data curation, Writing – original draft, Visualization. MGM: Methodology, Validation, Investigation. SG: Methodology, Validation, Formal analysis, Investigation, Data curation, Visualization. IZ: Validation, Investigation, Data curation. PR: Validation, Investigation, Data curation. YS: Supervision, Funding Acquisition, Project administration. MJ: Conceptualization, Supervision, Funding Acquisition, Project administration. JM: Conceptualization, Supervision, Funding Acquisition, Project administration. All the authors contributed in reviewing and editing the manuscript.

## Data availability

FIB-SEM and fluorescence light microscopy data of human BT-474 spheroids and patient-derived colorectal cancer organoids will be deposited on EMPIAR and Bioimage archive.

