## Supplementary Figures S1-S6 and captions to Movies 1-7 for "Light and electron microscopy continuum-resolution imaging of 3D cell cultures"

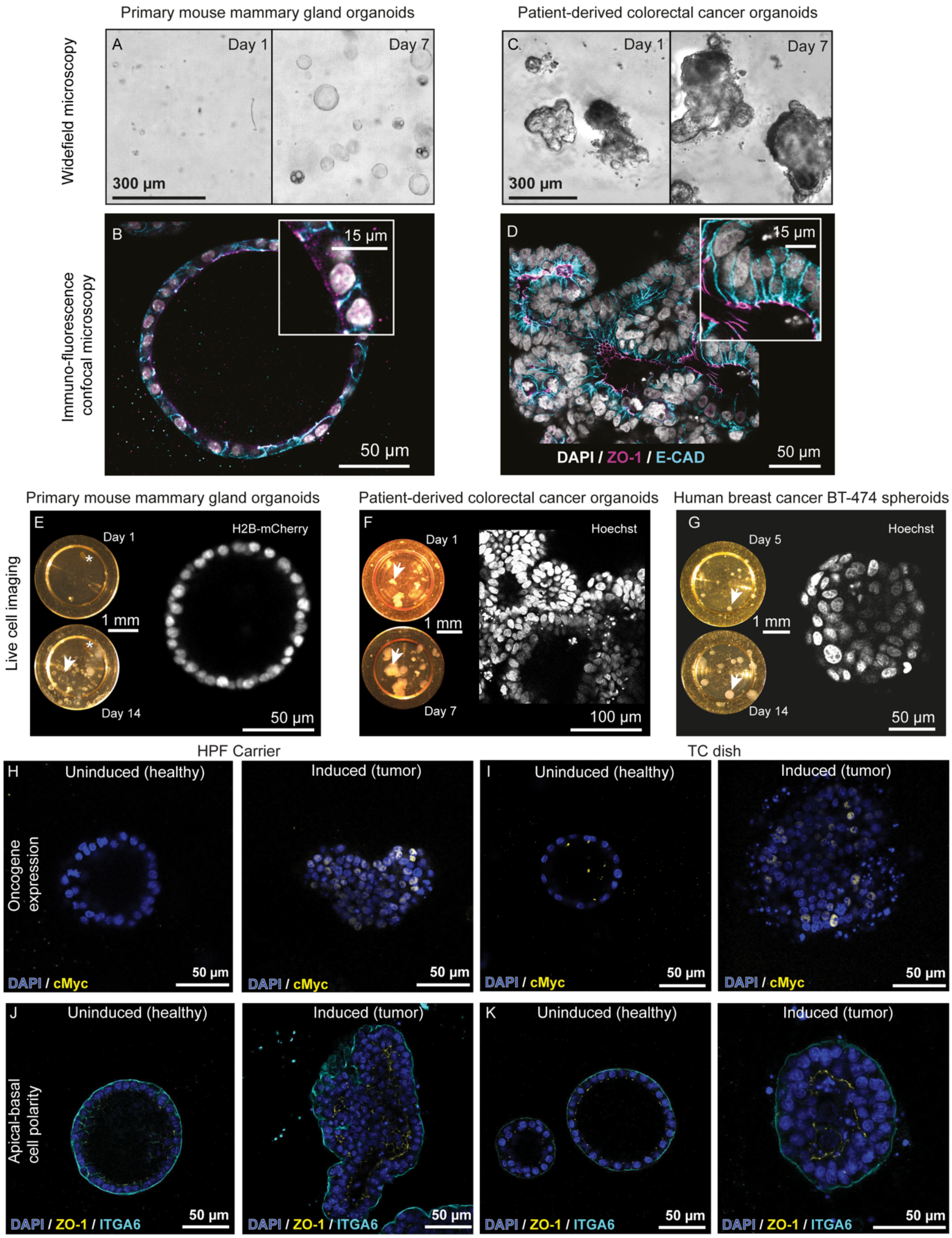


**Figure S1. Comparison of organoids growth in tissue culture dishes and HPF carriers.** **A**, growth of mouse mammary gland organoids in standard tissue culture (TC) dishes at the indicated time points. **B**, immunostaining in TC dishes of polarity markers: the cell-adhesion protein epithelial cadherin (E-CAD, cyan), tight junctions (ZO-1). Cell nuclei stained with DAPI (white). Inset: details of cell-cell interfaces. **C**, patient-derived colorectal cancer organoids growth at the indicated time in TC dishes. **D**, immunostaining in TC dishes of polarity markers

E-cadherin (E-CAD, cyan), tight junctions (ZO-1, magenta). Cell nuclei stained with DAPI (white). Inset: details of cell-cell interfaces. **E**, growth of mouse mammary gland organoids in HPF carrier at indicated time points. Asterisks indicate marks of the HPF carriers and arrowhead indicates a single organoid. Right: a confocal plane of an organoid at Day 14 expressing mCherry-tagged histone H2B. **F**, patient-derived colorectal cancer organoids growth in HPF carriers at the indicated time points of the culture. Arrowheads indicate growth of a single organoid. Right: a confocal plane shows an organoid at Day 7 with cell nuclei labelled with Hoechst-33342. **G**, human breast cancer spheroids growth at the indicated time points. Arrowheads indicate growth of a single organoid. Right: a confocal plane shows an organoid at Day 14 with cell nuclei labelled with Hoechst-33342. **H-I,** immunostaining of healthy and doxycycline-induced tumorigenic primary mouse luminal mammary gland organoids in HPF carriers (H) and TC dishes (I). Cell nuclei (DAPI, blue) and transgenic Myc oncogene expression (cMyc, yellow). **J-K**, immunostaining of healthy and tumorigenic organoids in HPF carriers (J) and TC dishes (K) showing expression of cell polarity markers: tight junctions (ZO-1, yellow) and integrin alpha-6 (ITGA6, green), an adhesion protein. Cell nuclei stained with DAPI (blue). Related to Figure 2.

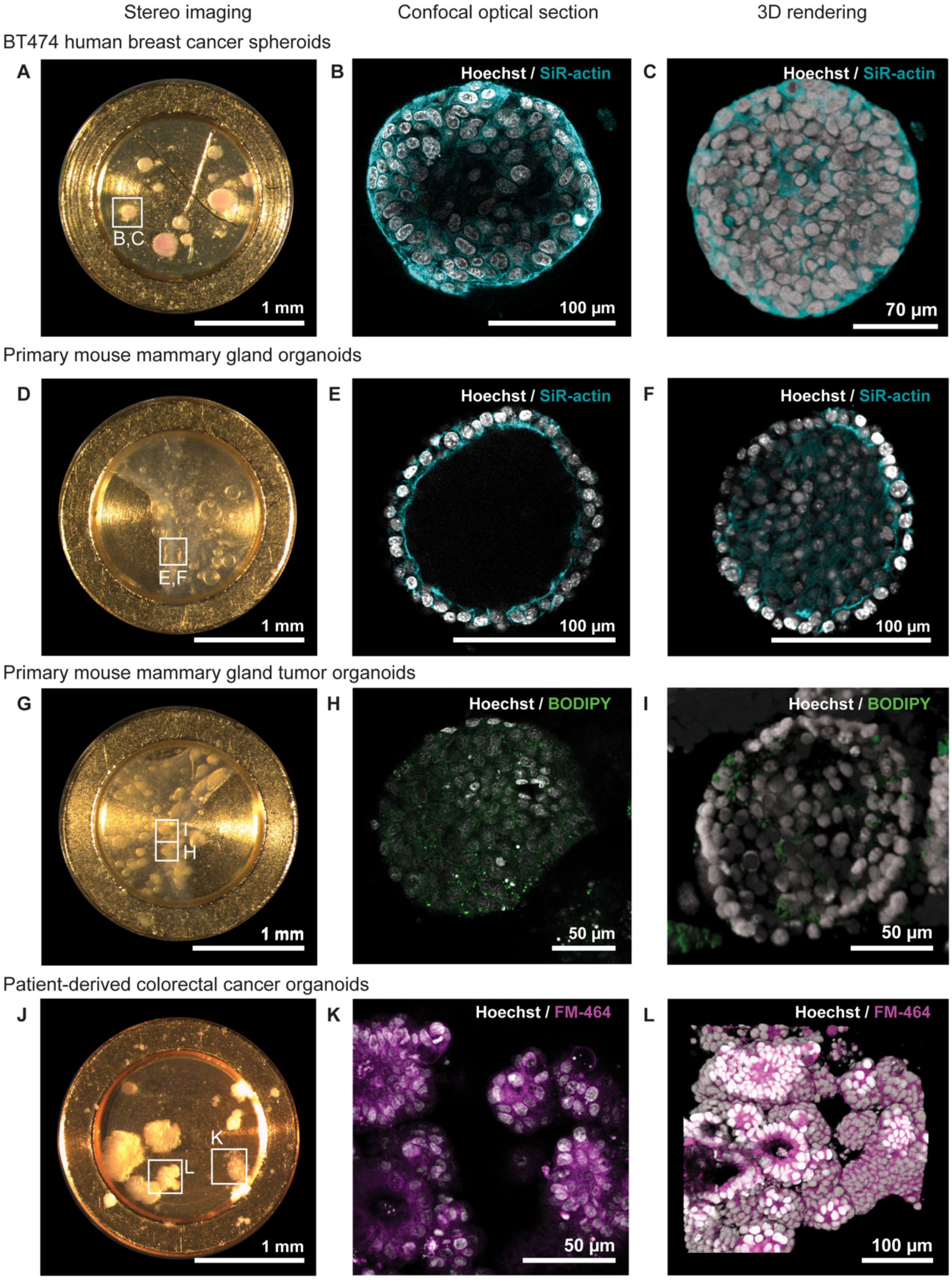


**Figure S2. Staining live cells from different 3D cell cultures with cell-permeable fluorescent dyes in HPF carriers.** Correlation of stereo microscopy (left, A, D, G, J), live-cell confocal fluorescence (center, B, E, H, K), and volume rendering (right, C, F, I, L) of four different 3D cell cultures. Organoids delineated by frames in the left column are enlarged in the right panels. Related to Figure 3.

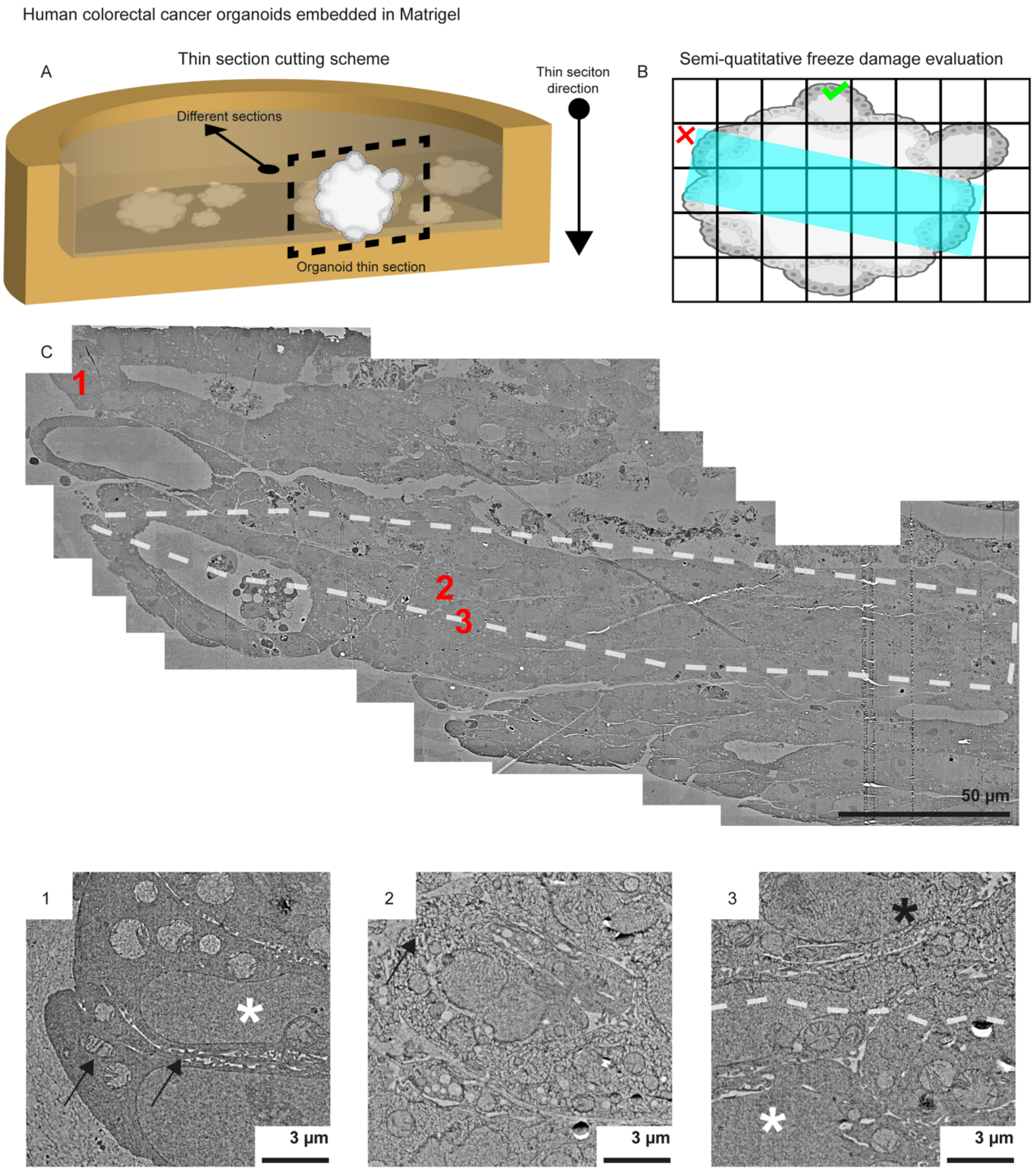


**Figure S3. Thin section TEM of human colorectal cancer organoids grown in HPF carriers for assessment of HPF efficiency. A**, a schematic illustrates that thin sections were cut perpendicular to the HPF carrier surface to achieve a cross-section of the bulk sample. Serial sections were collected advancing towards the center of the HPF carrier. **B**, the sections were imaged as a tiled montage (black mesh) and ultrastructural preservation was scored by counting tiles with evident freeze damage (red cross) versus tiles devoid of it (green tick). **C**, a tiled montage of a human colorectal cancer organoid. Freeze damage is confined in an area delineated by a white dashed line. Tiles indicated by red numbers are enlarged in the lower panels: (1) Cells exhibiting well-preserved nuclei (white asterisk), cell-cell interfaces, and mitochondria (black arrows). (2) Cells within the damaged area with evident ice segregation pattern in the cytosol (black arrow). (3) Cells at the interface (white dash) between damaged area (black asterisk) and preserved area (white asterisk). Images are displayed with positive contrast due to the different imaging of TEM compared to SEM. The contrast is inverted compared to FIB-SEM due to the different signal detection. Related to Figure 4.

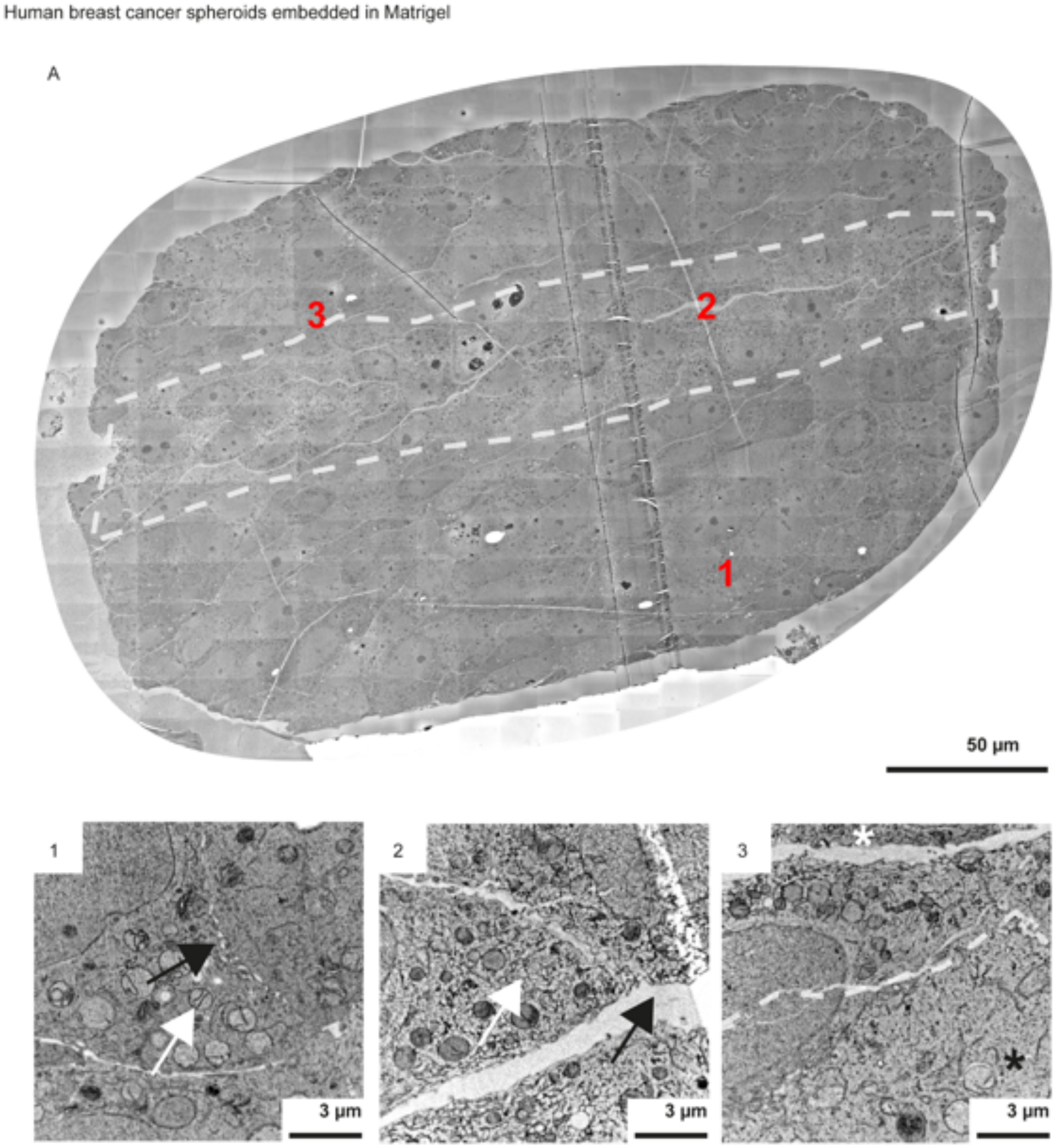


**Figure S4. Thin section TEM of human breast cancer spheroids grown in HPF carriers for assessment of HPF efficiency.** Top panel: (A) a tiled montage of a human breast cancer spheroid. Freeze damage is confined in an area delineated by the white dashed line and coincides with the center of the sample. Tiles indicated by red numbers are enlarged in the lower panels: (1) Cells outside the damaged area show details of well-preserved cell-cell interfaces (black arrows) and mitochondria (white arrows). (2) Cells within the damaged area show cytosol segregation (white arrow) and extensive cracks (black arrow). (3) Cells at the interface (white dash) between damaged (white asterisk) and preserved areas (black asterisk). Images are displayed with positive contrast due to the different imaging of TEM compared to SEM. The contrast is inverted compared to FIB-SEM due to the different signal detection. Related to Figure 4.

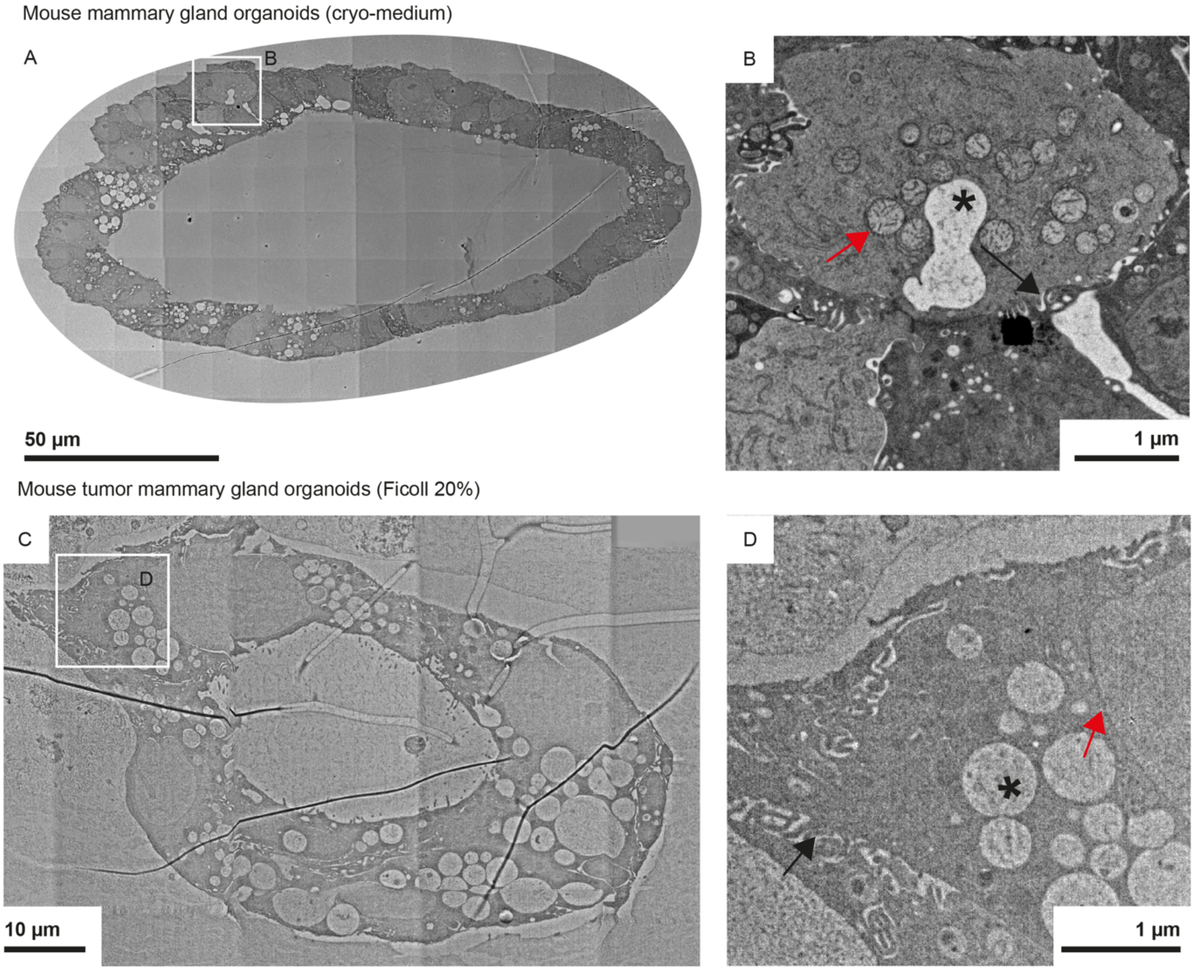


**Figure S5. Thin section TEM of mouse organoids grown in HPF carriers supplemented with cryo-protectants for one minute prior to freezing. A,** mouse organoids embedded in Matrigel and supplemented with Cellbanker 2 cryo-preserving medium (cryo-medium). **B**, enlarged view of the frame in A showing well preserved mitochondria (red arrows), cell-cell interfaces (black arrows), and vacuoles (black asterisk). **C**, mouse organoids embedded in Matrigel and supplemented with Ficoll (70.000 MW) 20%. **D**, details of well-preserved vacuoles (black asterisk), cell nucleus (red arrows), and cell-cell interfaces (black arrows). Images are displayed with positive contrast due to the different imaging of TEM compared to SEM. The contrast is inverted compared to FIB-SEM due to the different signal detection. Related to Figure 4.

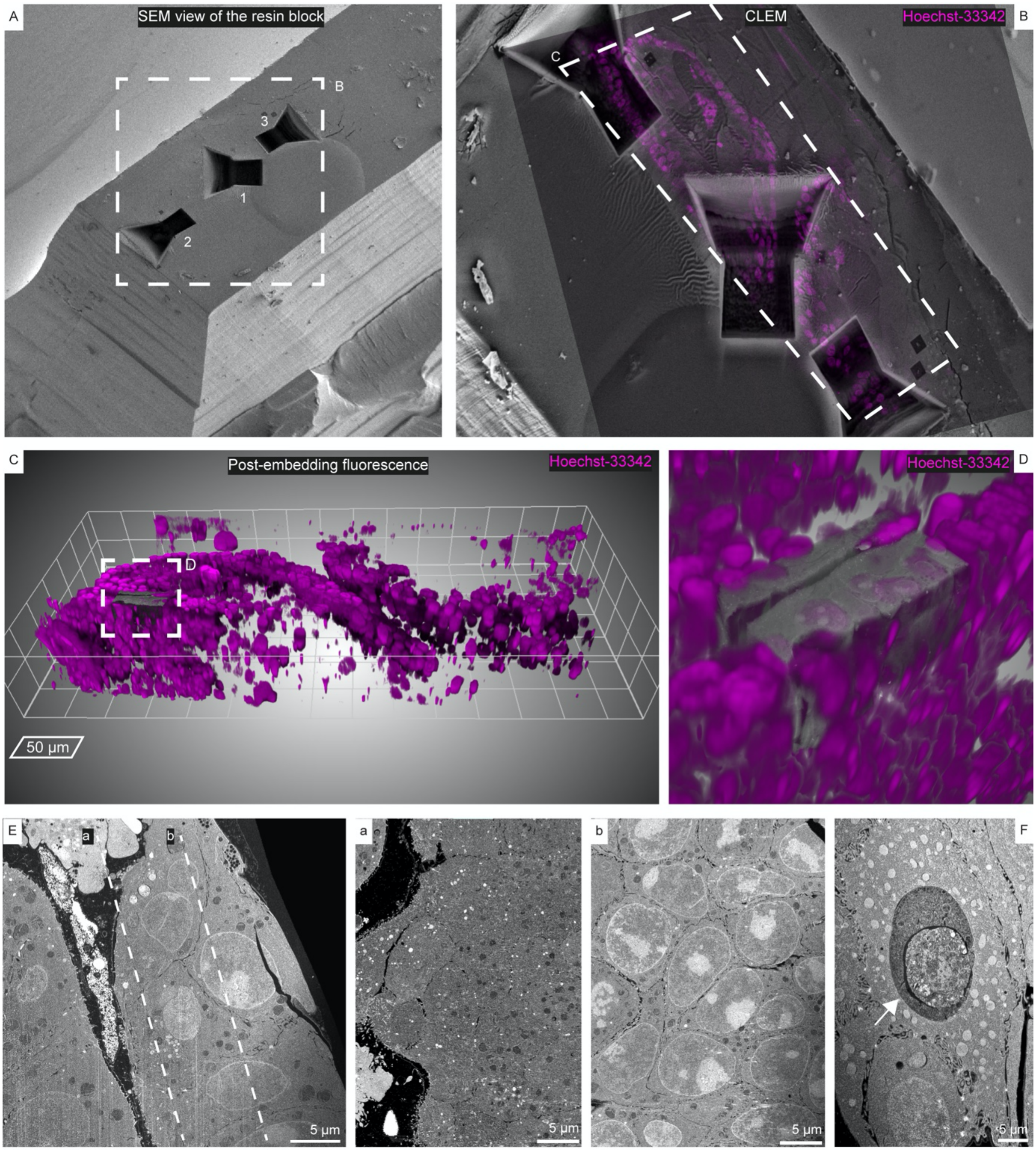


**Figure S6. Different cell packing morphologies within a single sample of patient-derived colorectal cancer organoid. A**, SEM view of a resin block containing Matrigel embedded patient-derived colorectal cancer organoids after HPF and freeze substitution. Based on the overlay of the post-embedding nuclear fluorescence (magenta, **B**), three areas showing different cell packing arrangements were target for FIB-SEM acquisitions (see also Movies 3, 4, 5). **C**-**D**, CLEM of one FIB-SEM acquisition (area 2) within the fluorescence volume of the complete patient-derived colorectal cancer organoid (mesh size 50 µm). **E**, single frame of a FIB-SEM acquisition of the target area 2 showing polarized epithelium. Two cross-section images in the volume were produced along the lines indicated by (a) and (b) and are shown on the right: intracellular morphologies with respect to the organoid lumen showing the apical (a) and basal (b) side of the cells. brighter signal likely corresponds to nucleoli. **F**, detail of an entotic cell (white arrow) from the acquisition in target are 1. Entosis, also known as “cell-in-cell structure”, involves the invasion or engulfment of one cell into the other in cancer tissues during epithelial to mesenchymal transition. Related to Figures 6 and 7.

**Movie 1. FIB-SEM volume imaging of cells within a human breast cancer spheroid.** Volume imaging of the single cells within the spheroid shown in Figure 5A. Labels highlight: mitochondria, vacuoles and nuclei. For display purposes the dataset was downscaled in x,y,z by half compared to the original FIB-SEM data acquired with voxels of 10x10x10 nm and the contrast adjusted by normalizing the local contrast in Fiji.

**Movie 2 . FIB-SEM volume imaging of a patient-derived colorectal cancer organoid with mixed arrangement of cell packing.** Related to acquisition position (3) in Figure S6A and Figure 7A (green). Labels highlight: nuclei, microvilli, lumina, one entotic cell and cell junctions. For display purposes the dataset was downscaled in x,y,z by half compared to the original FIB-SEM data acquired with voxels of 15x15x20 nm.

**Movie 3. FIB-SEM volume imaging of a patient-derived colorectal cancer organoid with monolayer epithelium cell packing.** Related to acquisition position (1) in Figure S6A and Figure 7A (red). Labels highlight: Lumen, nuclei, mitochondria, microvilli, condensed chromatin and cell junctions. For display purposes the dataset was downscaled in x,y,z by half compared to the original FIB-SEM data acquired with voxels of 15x15x20 nm.

**Movie 4. FIB-SEM volume imaging of a patient-derived colorectal cancer organoid with compact cell packing.** Related to acquisition position (2) in Figure S6A and Figure 7A (blue). Labels highlight: nuclei, mitochondria, cell junctions, lumina and microvilli. For display purposes the dataset was downscaled in x,y,z by half compared to the original FIB-SEM data acquired with voxels of 15x15x20 nm.

**Movie 5. FIB-SEM volume segmentation of a patient-derived colorectal cancer organoid with mixed arrangement of cell packing.** 2D rendering of the volume in Figure 7 (green) and Movie 2 showing the FIB-SEM data overlayed with labels of cell nuclei (light blue), entotic cell (gold), mitochondria (pink), actin bundles (magenta) and cell junctions (cyan). The segmentation is then rendered in 3D without FIB-SEM data overlayed.

**Movie 6. FIB-SEM volume segmentation of a patient-derived colorectal cancer organoid with monolayer epithelium cell arrangement.** 2D rendering of the data shown in Figure in Figure S6E, Figure 7 (red) and Movie 3 showing the FIB-SEM data overlayed with labels of cell nuclei (light blue), mitochondria (pink), actin bundles in microvilli (magenta) and cell junctions (cyan). The segmentation is then rendered in 3D without FIB-SEM data overlayed.

**Movie 7. FIB-SEM volume segmentation of a patient-derived colorectal cancer organoid with compact cell packing morphology.** 2D rendering of the data shown in Figure 7 (blue) and Movie 4 showing the FIB-SEM data, overlayed with labels of cell nuclei (light blue), mitochondria (pink), actin bundles in microvilli (magenta) and cell junctions (cyan). The segmentation is then rendered in 3D without FIB-SEM data overlayed.
